# Network Analysis Reveals Different Cellulose Degradation Strategies across *Trichoderma harzianum* Strains Associated with XYR1 and CRE1

**DOI:** 10.1101/2020.05.02.074344

**Authors:** Rafaela Rossi Rosolen, Alexandre Hild Aono, Déborah Aires Almeida, Jaire Alves Ferreira Filho, Maria Augusta Crivelente Horta, Anete Pereira de Souza

## Abstract

*Trichoderma harzianum*, whose gene expression is tightly controlled by the transcription factors (TFs) XYR1 and CRE1, is a potential candidate for hydrolytic enzyme production. Here, we performed a network analysis of *T. harzianum* IOC-3844 and *T. harzianum* CBMAI-0179 to explore how the regulation of these TFs varies between these strains. In addition, we explored the evolutionary relationships of XYR1 and CRE1 protein sequences among *Trichoderma* spp. The results of the *T. harzianum* strains were compared with those of *Trichoderma atroviride* CBMAI-0020, a mycoparasitic species. Although transcripts encoding carbohydrate-active enzymes (CAZymes), TFs, transporters, and proteins with unknown functions were coexpressed with *cre1* or *xyr1*, other proteins indirectly related to cellulose degradation were identified. The enriched GO terms describing the transcripts of these groups differed across all strains, and several metabolic pathways with high similarity between both regulators but strain-specific differences were identified. In addition, the CRE1 and XYR1 subnetworks presented different topology profiles in each strain, likely indicating differences in the influences of these regulators according to the fungi. The hubs of the *cre1* and *xyr1* groups included transcripts not yet characterized or described as being related to cellulose degradation. The first-neighbor analyses confirmed the results of the profile of the coexpressed transcripts in *cre1* and *xyr1*. The analyses of the shortest paths revealed that CAZymes upregulated under cellulose degradation conditions are most closely related to both regulators, and new targets between such signaling pathways were discovered. Although the evaluated *T. harzianum* strains are phylogenetically close and their amino acid sequences related to XYR1 and CRE1 are very similar, the set of transcripts related to *xyr1* and *cre1* differed, suggesting that each *T. harzianum* strain used a specific regulation strategy for cellulose degradation. More interestingly, our findings may suggest that XYR1 and CRE1 indirectly regulate genes encoding proteins related to cellulose degradation in the evaluated *T. harzianum* strains. An improved understanding of the basic biology of fungi during the cellulose degradation process can contribute to the use of their enzymes in several biotechnological applications and pave the way for further studies on the differences across strains of the same species.

## 1 Introduction

Lignocellulosic biomass is a complex recalcitrant structure that requires a consortium of carbohydrate-active enzymes (CAZymes) for its complete depolymerization. Due to their unique ability to secrete these proteins efficiently, filamentous fungi, such as *Trichoderma* spp. and *Aspergillus* spp., are widely explored for the industrial production of CAZymes (Bischof, et al. 2016; de Assis, et al. 2015). In the genus *Trichoderma*, *Trichoderma reesei* is the primary fungal industrial source of cellulases and hemicellulases (Martinez, et al. 2008), while *Trichoderma harzianum* and *Trichoderma atroviride* have been widely explored by examining their biocontrol capacity against plant pathogenic fungi (Medeiros, et al. 2017; Saravanakumar, et al. 2017). However, hydrolytic enzymes from *T. harzianum* strains have demonstrated great potential in the conversion of lignocellulosic biomass into fermentable sugars (Delabona, et al. 2020; Zhang, et al. 2020).

The production of CAZymes in filamentous fungi is controlled at the transcriptional level by several positive and negative transcription factors (TFs) (Benocci, et al. 2017). In *T. reesei*, the Zn_2_Cys_6_-type TF xylanase regulator 1 (XYR1) is described as the most important activator of cellulase and xylanase gene expression (Stricker, et al. 2006). During growth on an induction carbon source, XYR1 was shown to be synthesized *de novo* and degraded at the end of induction (Lichius, et al. 2014). Although XYR1 orthologs are present in almost all filamentous ascomycete fungi, the molecular mechanisms triggered by this regulator depend on the species (Klaubauf, et al. 2014). In *T. atroviride*, the induction of genes that encode cell wall-degrading enzymes considered relevant for mycoparasitism, such as *axe1* and *swo1*, is influenced by XYR1 (Reithner, et al. 2014). In *T. harzianum*, the overexpression of *xyr1* increases the levels of the reducing sugars released in a shorter time during saccharification (Delabona, et al. 2017). Overall, in the genus *Trichoderma*, XYR1 evolved by vertical gene transfer (Druzhinina, et al. 2018).

In the presence of easily metabolizable carbon sources, such as glucose, the expression of *xyr1* and genes encoding lignocellulose-degrading enzymes is repressed by carbon catabolite repression (CCR), which is regulated by the C_2_H_2-_type TF carbon catabolite repressor 1 (CRE1) (Alazi and Ram 2018; Mach-Aigner, et al. 2008; Strauss, et al. 1995). Upon glucose depletion, the concentration of CRE1 in the nucleus rapidly decreases, and CRE1 is recycled into the cytoplasm (Lichius, et al. 2014). Furthermore, the phosphorylation of CRE1 plays an essential role in signal transduction to achieve CCR (Han, et al. 2020; Horta, et al. 2019). CRE1 is the only conserved TF throughout the fungal kingdom, suggesting a conserved mechanism for CCR in fungi (Adnan, et al. 2017). Interestingly, the effect of *cre1* deletion on the gene expression profile of its targets varies among species. In *T. reesei* RUT-C30, the deletion of a full *cre1* can lead to pleiotropic effects and strong growth impairment (Mello-de-Sousa, et al. 2014; Portnoy, et al. 2011), whereas in *T. harzianum* P49P11, it can increase the expression levels of CAZyme genes and *xyr1* (Delabona, et al. 2021).

Although *T. harzianum* strains have shown high cellulolytic activity (Horta, et al. 2018; Li, et al. 2020b), most studies investigating this species have focused on biological control (Maruyama, et al. 2020; Yan and Khan 2021). Thus, the regulatory mechanisms underlying hydrolytic enzyme production by these fungi are still poorly explored at the transcriptional level (Delabona, et al. 2017, 2020, 2021). In addition, given the high degree of genetic variation observed within the genus *Trichoderma* (Kubicek, et al. 2019) along with the complex speciation observed in the *T. harzianum* species (Druzhinina, et al. 2010), it is necessary to explore how the main regulators XYR1 and CRE1 behave across *T. harzianum* strains.

Previously, the transcriptomes of *T. harzianum* IOC-3844 (Th3844), *T. harzianum* CBMAI-0179 (Th0179), and *T. atroviride* CBMAI-0020 (Ta0020) were investigated under cellulose degradation conditions (Almeida, et al. 2021). Different types of enzymatic profiles were reported, and both *T. harzianum* strains had higher cellulase activity than *T. atroviride*. Using this dataset, we aimed to investigate how the regulation of the TFs CRE1 and XYR1 varies among *T. harzianum* strains. Based on the assumption that coexpressed genes tend to share similar expression patterns and that they could be coregulated by the same elements, we modeled a network of Th3844, Th0179, and Ta0020 using a weighted correlation network analysis (WGCNA) (Langfelder and Horvath 2008). The last strain, which is distantly related to *T. reesei* (Druzhinina, et al. 2006) and represents a well-defined phylogenetic species (Dodd, et al. 2003), was used to assess the differences across *Trichoderma* species. In addition, phylogenetic analyses of XYR1 and CRE1 protein sequences were performed to clarify the evolutionary relationships of these regulators among the evaluated strains.

In this study, we identified and compared modules, hub genes, and metabolic pathways associated with CRE1 and XYR1 under cellulose degradation conditions. To deeply investigate their regulatory activities, we also performed first neighbor and shortest-path network analyses. Although the evaluated *T. harzianum* strains are phylogenetically close, by comparing their coexpressed transcript profiles, functional diversity was observed. This difference was accentuated when associating the results of Th0179 and Th3844 with those of Ta0020. Thus, we observed a specific transcriptional pattern related to CRE1 and XYR1 in each strain. Our study could contribute to improving our understanding of the regulation of cellulose degradation in *T. harzianum* and paves the way for further studies evaluating differences across strains within the same species. Addressing these questions by investigating the genetic regulatory mechanisms involved in cellulose degradation is important for enhancing both our basic understanding and biotechnological industrial applications of fungal abilities.

## 2 Materials and methods

### 2.1 Fungal strains, culture conditions and transcription profiling

The species were obtained from the Brazilian Collection of Environment and Industry Microorganisms (CBMAI) located in the Chemical, Biological, and Agricultural Pluridisciplinary Research Center (CPQBA) of the University of Campinas (UNICAMP), Brazil. The identity of the *Trichoderma* isolates was authenticated by CBMAI based on phylogenetic studies of their internal transcribed spacer (ITS) region and translational elongation factor 1 (*tef1*) marker gene. The *T. harzianum* CBMAI-0179 (Th0179), *T. harzianum* IOC-3844 (Th3844), and *T. atroviride* CBMAI-0020 (Ta0020) strains were cultivated in simple carbon sources, including cellulose and glucose, to induce a clear genetic response. Details on the culture conditions and transcription profiling of the described experiments are provided in Supplementary Material 1, while the profile of enzymatic activities (cellulase, beta-glucosidase, and xylanase), the total protein content, and the constitution of the secreted proteome has been published previously (Almeida, et al. 2020; Horta, et al. 2018).

### 2.2 Phylogenetic analyses

The ITS nucleotide sequences of *Trichoderma* spp. were retrieved from the NCBI database (https://www.ncbi.nlm.nih.gov/). Additionally, ITS nucleotide sequences amplified from the genomic DNA of Th3844, Th0179, and Ta0020 via PCR were kindly provided by the CBMAI and included in the phylogenetic analysis. The ITS region has been found to be among the markers with the highest probability of correctly identifying a very broad group of fungi (Schoch, et al. 2012). *T. harzianum* T6776 was used as a reference genome to retrieve the CRE1 and XYR1 protein sequences belonging to *Trichoderma* spp. Additionally, the CRE1 and XYR1 sequences were obtained from the *T. harzianum* T6776 (Baroncelli, et al. 2015) reference genome of Th3844 and Th0179 and the *T. atroviride* IMI206040 (Kubicek, et al. 2011) genome of Ta0020 and used as references for the TF consensus sequences for RNA-Seq read mapping. These sequences were included in the phylogenetic analyses.

The multiple sequence alignment was performed using ClustalW (Thompson, et al. 1994), and a phylogenetic tree was created using Molecular Evolutionary Genetics Analysis (MEGA) software v7.0 (Kumar, et al. 2016). The maximum likelihood (ML) (Jones, et al. 1992) method of inference was used based on (I) a Kimura two-parameter (K2P) model (Kimura 1980), (II) a Dayhoff model with frequencies (F^+^), and (III) a Jones-Taylor-Thornton (JTT) model for ITS, CRE1, and XYR1, respectively. We used 1,000 bootstrap replicates (Felsenstein 1985) in each analysis. The trees were visualized and edited using Interactive Tree of Life (iTOL) v6 (https://itol.embl.de/).

### 2.3 Weighted gene coexpression network analysis

The gene coexpression networks of Th3844, Th0179, and Ta0020 were modeled using transcripts per million (TPM) value data, which were validated through qPCR and described by Almeida *et al.* (2021), of three biological replicates with the R (R Core Team 2018) WGCNA package. Transcripts showing null values for most replicates under different experimental conditions were excluded. The network was assembled by calculating the Pearson’s correlation coefficient of each pair of genes. A soft power β was chosen for each network using *pickSoftThreshold* to fit the signed network to a scale-free topology. Then, an adjacency matrix in which the nodes correspond to transcripts and the edges correspond to the strength of their connection was obtained. To obtain a dissimilarity matrix, we built a topological overlap matrix (TOM) as implemented in the package.

To identify groups of transcripts densely connected in the gene coexpression network, we applied simple hierarchical clustering to the dissimilarity matrix. From the acquired dendrogram, the *dynamicTreeCut* package (Langfelder, et al. 2008) was used to obtain the ideal number of groups and the respective categorization of the transcripts. According to the functional annotation performed by Almeida, et al. (2021), groups containing the desired TFs XYR1 and CRE1 were identified and named the *xyr1* and *cre1* groups, respectively.

### 2.4 Functional annotation of transcripts in the xyr1 and cre1 groups

We functionally annotated the transcripts in the *xyr1* and *cre1* groups. By conducting a Fisher’s exact test to extract the overrepresented terms (p-value < 0.05), all identified Gene Ontology (GO) (Ashburner, et al. 2000) categories were used to identify enriched GO terms with the topGO package in R (Alexa and Rahnenfuhrer 2021). To visualize the possible correlated enriched categories in the dataset caused by cellulose degradation, we created a treemap using the REVIGO tool (Supek, et al. 2011). Then, the metabolic pathways related to the Kyoto Encyclopedia of Genes and Genomes (KEGG) (Kanehisa and Goto 2000) Orthology (KO) identified in *T. reesei* were selected due the high number of annotations correlated with this species in the KEGG database. To identify the pathways related to the enzymes identified as belonging to different groups, we used the Python 3 programming language (Sanner 1999) along with the BioPython library (Cock, et al. 2009). The automatic annotation was manually revised using the UniProt databases (UniProt Consortium 2019).

### 2.5 Assessing the groups’ network topologies

To provide a visual network characterization, we modeled additional gene coexpression networks of all strains using the highest reciprocal rank (HRR) approach (Mutwil, et al. 2010). Using an R Pearson correlation coefficient threshold of 0.8, we assessed the 30 strongest edges, which were coded as ⍰ (top 10), 1/15 (top 20), and 1/30 (top 30). To visualize the behavior of the groups modeled by WGCNA in the HRR networks, we used Cytoscape software v3.7.0 (Shannon, et al. 2003). Using the HRR methodology, we were able to infer the possible biological correlations depending on the network topology. First, by performing a topological analysis based on the degree distribution, we identified and compared the hub nodes in the *cre1* and *xyr1* groups among all evaluated strains. Second, to identify the more indirect associations of both TFs studied, we considered HRR global networks to evaluate the first neighbors of the *xyr1* and *cre1* transcripts. Finally, because XYR1 and CRE1 are described as regulatory proteins in CAZyme gene expression, we were also interested in evaluating the minimum pathway between these regulatory proteins and these hydrolytic enzymes. Based on the classification proposed by Almeida, et al. (2021), we selected CAZymes with higher expression levels under cellulose growth conditions (upregulated transcripts) that were present in both Th0179 and Th3844. To obtain the corresponding homologs of Ta0020, BLASTp was performed using *T. atroviride* IMI206040 as the reference genome. The numbers of the shortest paths between both TFs, namely, CRE1 and XYR1, and the selected CAZymes were identified using the Pesca v. 3.0 plugin (Scardoni, et al. 2016) in Cystoscope. The results were visualized using the R package pheatmap (Kolde 2019).

## 3 Results

### 3.1 Molecular phylogeny of the evaluated *T. harzianum* strains

Although Kubicek, et al. (2019) deeply investigated the phylogenetic relationships in the genus *Trichoderma*, no results were reported for Th3844, Th0179, and Ta0020. Here, we modeled a phylogenetic tree based on the ITS sequence of 14 *Trichoderma spp*., including those of our study strains (Supplementary Material 1: Supplementary Figure 1). According to the results, high genetic proximity between Th3844 and Th0179 was observed. In contrast, both strains were phylogenetically distant from Ta0020, which grouped with other *T. atroviride* strains. *Neurospora crassa* and *Fusarium oxysporum* were used as outgroups.

Additionally, to represent the evolutionary relationships of CRE1 and XYR1 among Th3844, Th0179, and Ta0020, a phylogenetic analysis was performed while considering their amino acid sequences (Figure 1). The phylogenetic trees were modeled based on 26 and 21 protein sequences related to CRE1 and XYR1, respectively, in the *Trichoderma* genus, including those of our studied strains. *Fusarium* spp. was used as an outgroup. The NCBI accession number of the sequences used to model the phylogenetic trees is available in Supplementary Material 2: Supplementary Table 2.

**Figure 1.**
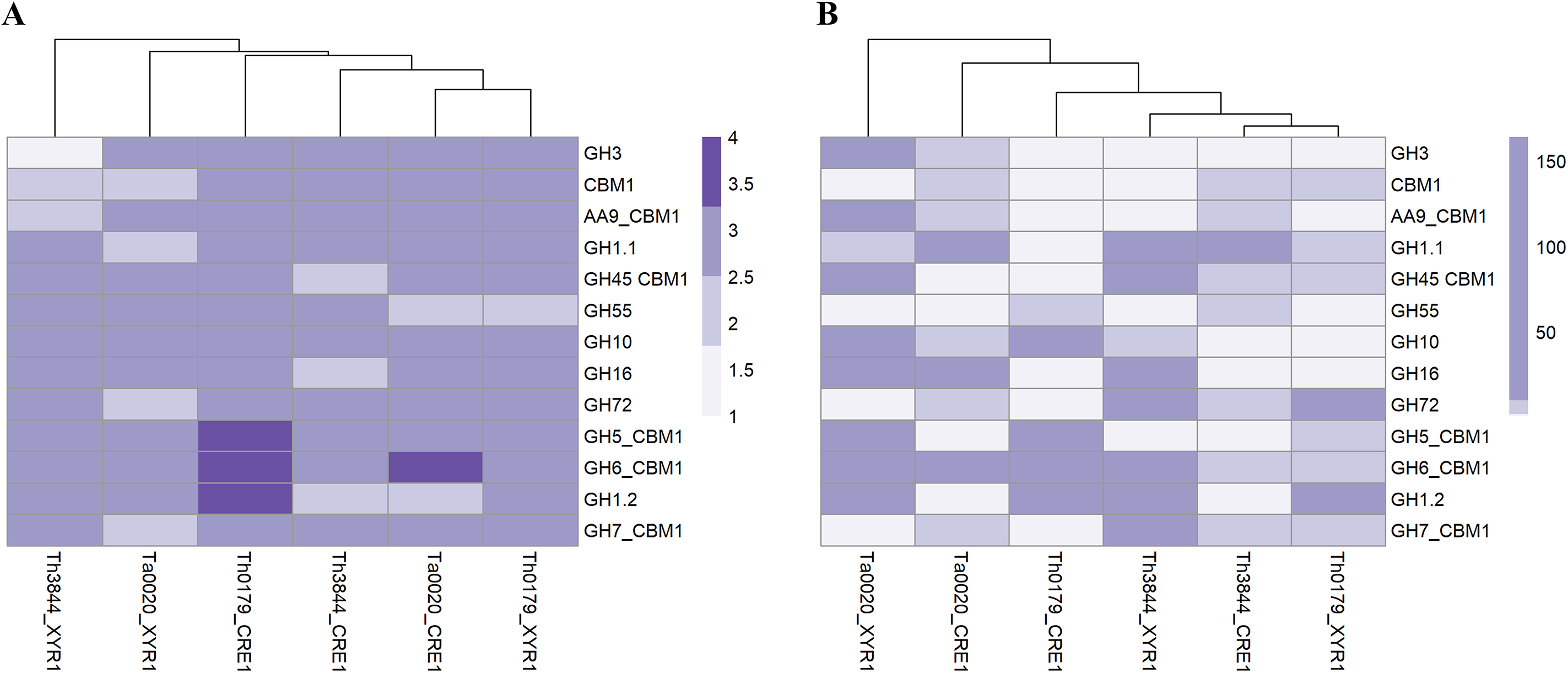
Molecular phylogenies of CRE1 and XYR1 in *Trichoderma* spp. The 26 and 21 complete protein sequences related to CRE1 (**A**) and XYR1 (**B**), respectively, were used to infer the phylogenetic relationships of the studied strains in the genus *Trichoderma*. Close genetic proximity between Th3844 and Th0179 was observed in the CRE1 phylogenetic tree, while Ta0020 shared a closer affinity with other *T. atroviride* strains. In the XYR1 phylogenetic tree, Th3844, Th0179, and Ta0020 were closest to *T. harzianum* CBS 226.95, *T. harzianum* T6776, and *T. atroviride* IMI206040, respectively. *Fusarium* spp. was used as an outgroup. The bootstrap value is represented in each node. The phylogenetic trees were constructed using MEGA7 software and edited using the iTOL program.

The CRE1 phylogenetic tree indicated a close genetic proximity between Th3844 and Th0179 (with bootstrap support of 67%) (Figure 1A). These results were supported by the alignment of both protein sequences, which showed high conservation in the alignment of their amino acid sequences (with a percent identity of 99.76%) (Supplementary Material 1: Supplementary Figure 2). In contrast, the CRE1 protein sequence of Ta0020 shared closer affinity with two other *T. atroviride* strains (T23 and IMI206040), with bootstrap support of 99% (Figure 1A). Furthermore, the C_2_H_2_ domain is highly conserved in the Th3844, Th0179, and Ta0020 sequences of CRE1, with only one amino acid change from *T. atroviride* compared to both *T. harzianum* strains (Supplementary Material 1: Supplementary Figure 2).

In the XYR1 phylogenetic tree, Th3844, Th0179, and Ta0020 were closest to *T. harzianum* CBS 226.95 (with bootstrap support of 97%), *T. harzianum* T6776 (with bootstrap support of 51%), and *T. atroviride* IMI206040 (with bootstrap support of 88%), respectively (Figure 1B). Compared to the results of CRE1, the alignment of their amino acid sequences showed a lower percentage of identity, supporting the phylogenetic results observed (Supplementary Material 1: Supplementary Figure 2). In addition, the multiple alignments of the XYR1 protein sequence of Th3844, Th0179, and Ta0020 showed reasonable conservation of the Zn2Cys6 and fungal-specific TF domains (Supplementary Material 1: Supplementary Figure 2).

### 3.2 Weighted gene coexpression network analysis

Recently, the transcriptomes of Th3844, Th0179, and Ta0020 grown on crystalline cellulose and glucose after a time course of 96 h were investigated (Almeida, et al. 2021). By using these transcriptome data and applying the described filters in the WGCNA package, we obtained (I) 11,050 transcripts (Th3844), (II) 11,105 transcripts (Th0179), and (III) 11,021 transcripts (Ta0020) (Supplementary Material 3: Supplementary Table 3). Based on these transcripts, we calculated Pearson’s correlation matrices. Then, different soft power β values were chosen (48 for Th3844, 8 for Th0179, and 27 for Ta0020) to obtain the scale-free topology, reaching fit indexes of (I) 0.8 for Th3844 (mean connectivity of 40), (II) 0.9 for Th0179 (mean connectivity of 842), and (III) 0.85 for Ta0020 (mean connectivity of 406). Then, the networks were partitioned into manageable groups to explore the putative coregulatory relationships. In total, we identified 87 groups in Th3844, 75 groups in Th0179, and 100 groups in Ta0020 (Supplementary Material 4: Supplementary Table 4). Transcripts with expression patterns correlated with XYR1 and CRE1 were identified in the Th3844, Th0179, and Ta0020 coexpression networks and grouped (described in Supplementary Material 5: Supplementary Table 5). Among the strains, Ta0020 presented the highest number of transcripts coexpressed with *cre1* and the lowest number of transcripts coexpressed with *xyr1*; the other strains showed the opposite profile.

### 3.3 Functional characterization of the transcripts in the *xyr1* and *cre1* groups

We performed an enrichment analysis of the transcripts in each *cre1* and *xyr1* group of all evaluated strains using the GO categories (Supplementary Material 1: Supplementary Figures 3-5, respectively). The *cre1* group of Th3844, Th0179, and Ta0020 presented 50, 38, and 50 enriched GO terms, respectively, while in the *xyr1* group, 34, 86, and 50 enriched GO terms were found in Th3844, Th0179, and Ta0020, respectively. Overall, the GO terms describing the transcripts of these groups were diverse across all strains, indicating a high degree of differences among the biological processes related to XYR1 and CRE1.

For example, the enrichment analyses of Ta0020 indicated that the carbohydrate metabolic process might be directly affected by CRE1 (high percentage of enriched GOs). Furthermore, GO terms related to fungal growth and nucleoside transmembrane transport were observed. In contrast, in Th0179 and Th3844, organic substance metabolic processes and regulation processes were pronounced, respectively. Interestingly, response to light stimulus was an enriched GO term in the *cre1* group of Th3844, and fungal-type cell wall organization was an enriched GO term in Th0179. In the XYR1 group, Ta0020 also presented carbohydrate metabolic process as an enriched GO term. However, few genes corresponding to this term were observed, while terms related to the regulation of DNA transcription were pronounced. Interestingly, response to external stimulus and oxidation-reduction process were notable terms in Th0179 and Th3844, respectively. Furthermore, our results suggest that the regulation of biological processes was a common enriched term in all the evaluated strains, with some particularities across the *T. harzianum* strains, including regulation of cell population proliferation (Th0179) and mRNA metabolic process (Th3844).

The KO functional annotation of the transcripts encoding enzymes and unknown proteins with enzymatic activity in the *cre1* and *xyr1* groups was also investigated. In the 14 pathway classes observed in each TF, 13 were shared between the two regulators (Figure 2). Only two pathway classes, i.e., metabolism of other amino acids and lipid metabolism, were exclusive to CRE1 and XYR1, respectively. However, the number of identified enzymes in a given pathway class differed in each strain. This difference stresses the diversity of proteins with several functions, which could result in exclusive enzymatic performance in cellulose degradation according to the strain.

**Figure 2.**
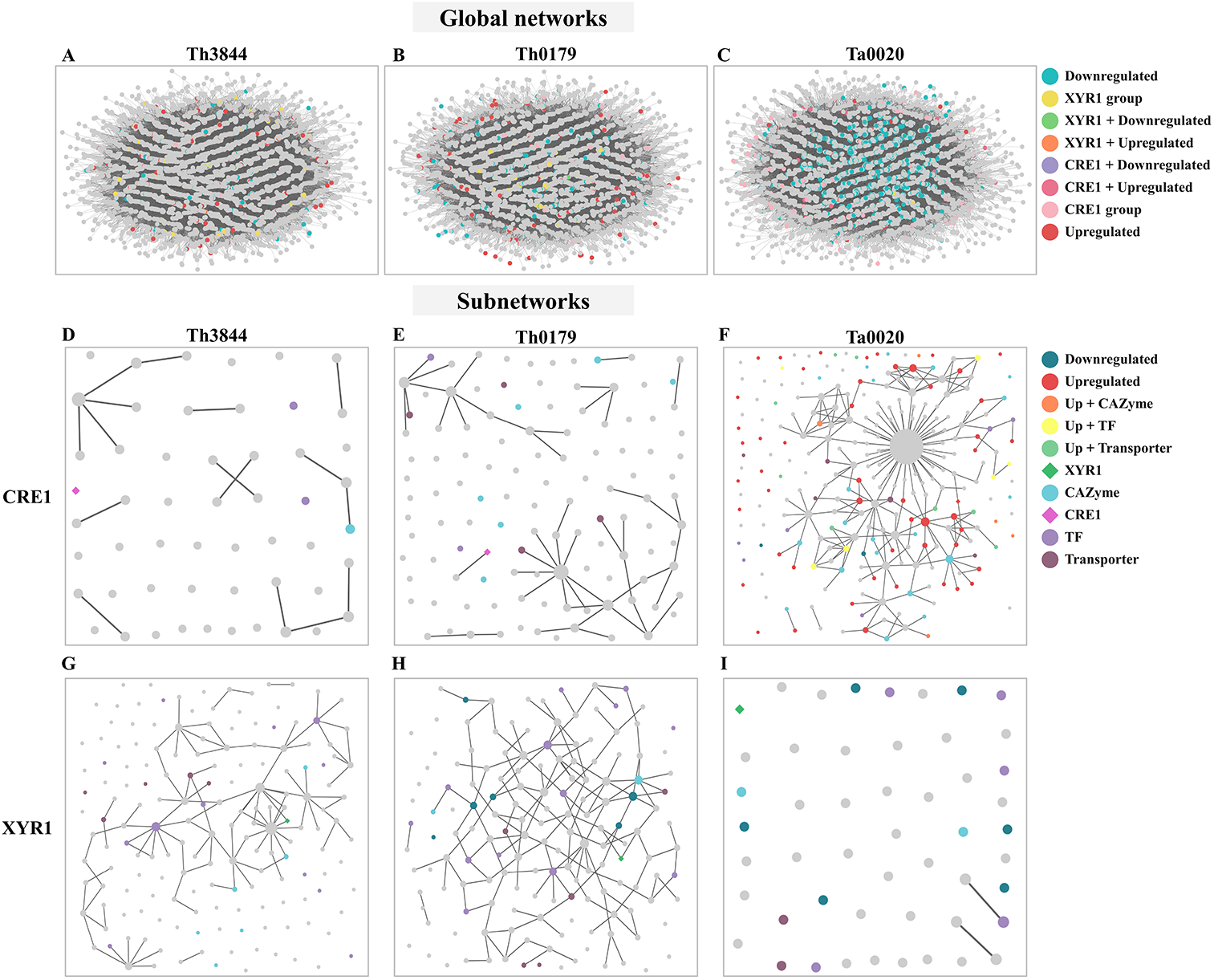
KO functional classification of the transcripts identified in the *cre1* and *xyr1* groups. The sequences related to the enzymes coexpressed with *cre1* (**A**) and *xyr1* (**B**) were annotated according to the main KO functions of Th3844, Th0179, and Ta0020. For all evaluated strains, 14 pathway classes were observed in each TF, and 13 classes were shared between the two regulators. However, the number of identified enzymes in a given pathway class differed in each strain. Each strain is represented by a different color. Ta0020: *T. atroviride* CBMAI-0020; Th0179: *T. harzianum* CBMAI-0179; Th3844: *T. harzianum* IOC-3844; KO: Kyoto Encyclopedia of Genes and Genomes Orthology.

Next, we summarize some important aspects of the identified metabolic pathways. For example, Ta0020 showed a greater number of enriched pathway classes in the *cre1* group, while both *T. harzianum* strains presented the opposite profile, with more enzymatic activity pathways enriched in the *xyr1* group. Carbohydrate metabolism was a pathway class enriched in both *T. harzianum* strains in the *xyr1* group, while Th0179 and Ta0020 presented such a profile in the *cre1* group. In the *xyr1* group, nucleotide metabolism and metabolism of cofactors and vitamins were enriched terms in Th0179, while xenobiotic degradation and metabolism of terpenoids and polyketides were enriched terms exclusively in Ta0020. The pathway classes related to secondary metabolite compounds were also enriched in all evaluated strains in the *cre1* and *xyr1* groups, and Ta0020 presented the highest number of enzymes related to this term in the *cre1* group.

In this study, network analyses were conducted to investigate how the molecular basis of XYR1 and CRE1 for cellulose-degrading enzyme production varies among *T. harzianum* strains. Thus, after analyzing the functional composition of the transcripts in the *cre1* and *xyr1* groups of all evaluated strains, particular attention was given to CAZymes, TFs, and transporters (Figure 3). We observed that in the *cre1* group, Ta0020 showed the highest number of transcripts encoding CAZymes, TFs, and transporters, while in the *xyr1* group, both *T. harzianum* strains presented a great number of coexpressed transcripts encoding TFs, followed by those encoding transporters. Furthermore, Th3844 showed a high number of CAZymes in the *xyr1* group.

**Figure 3.**
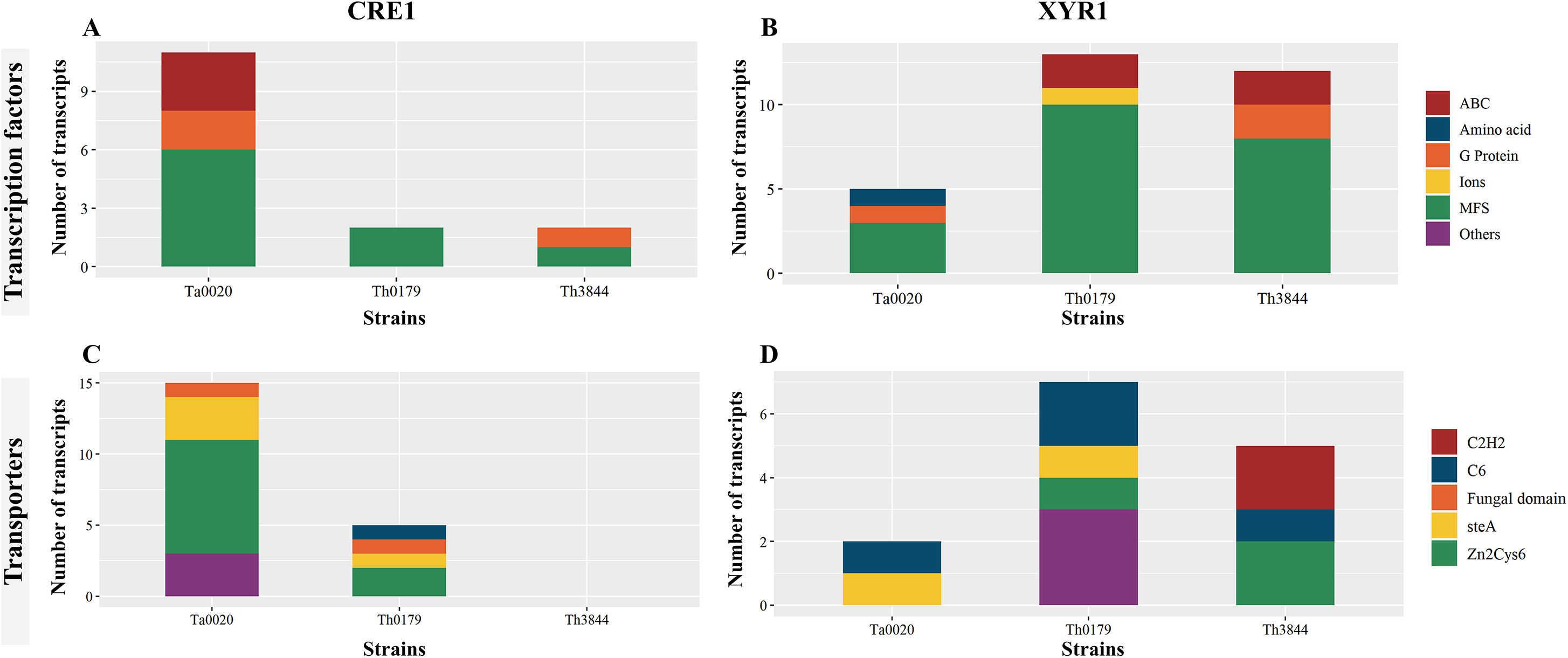
Comparisons of TFs, transporters, and CAZymes among strains and groups. Number of transcripts encoding TFs, transporters, and CAZymes in Ta0020, Th0179, and Th3844 distributed among the *cre1* (**A**) and *xyr1* (**B**) groups. The number of transcripts encoding these proteins associated with *cre1* was more accentuated in Ta0020 than those observed for the other strains, while in the *xyr1* group, both *T. harzianum* strains presented a great number of coexpressed transcripts encoding TFs, followed by those encoding transporters. Th3844 also showed a high number of CAZymes in the *xyr1* group. TFs: transcription factors; CAZymes: carbohydrate-active enzymes. Ta0020: *T. atroviride* CBMAI-0020; Th0179: *T. harzianum* CBMAI-0179; Th3844: *T. harzianum* IOC-3844.

We identified the main CAZyme classes, including glycoside hydrolases (GHs), carbohydrate esterases (CEs), glycosyltransferases (GTs), and polysaccharide lyases (PLs), coexpressed with *xyr1* and *cre1* transcripts. To determine the similarities and differences across the strains, their CAZyme profiles were compared (Figure 4). In the *xyr1* and *cre1* groups, the GH family, which encompasses enzymes that hydrolyze glycosidic bonds, was identified in all the evaluated strains with different numbers of transcripts (Figure 4A and B). In the *cre1* group, Ta0020 presented the highest number of classified transcripts of the GH family, followed by Th0179 and Th3844. In the *xyr1* group, Th3844 presented a higher number of GHs than Th0179 and Ta0020. Furthermore, transcripts encoding CEs, which hydrolyze ester bonds, were associated with the *cre1* transcript in Ta0020 and Th0179, whereas transcripts of the PL family, which cleave bonds in uronic acid-containing polysaccharide chains, were coexpressed with the *xyr1* transcript in only Th3844. In addition, transcripts of the GT family, which synthesizes glycosidic bonds from phosphate-activated sugar donors, were present in the *cre1* group of Th0179 and the *xyr1* groups of Th3844 and Ta0020. We also investigated the quantification of each CAZyme family in the *cre1* and *xyr1* groups of all evaluated strains (Figure 4C and D). Overall, Ta0020 presented a high number of GHs belonging to family 18, coexpressed with the *cre1* transcript, while Th0179 exhibited a significant amount of GT90 (Figure 4C). Different CAZyme profiles were observed in the *xyr1* groups, and Th3844 showed the highest diversity of CAZyme families (Figure 4D).

**Figure 4.**
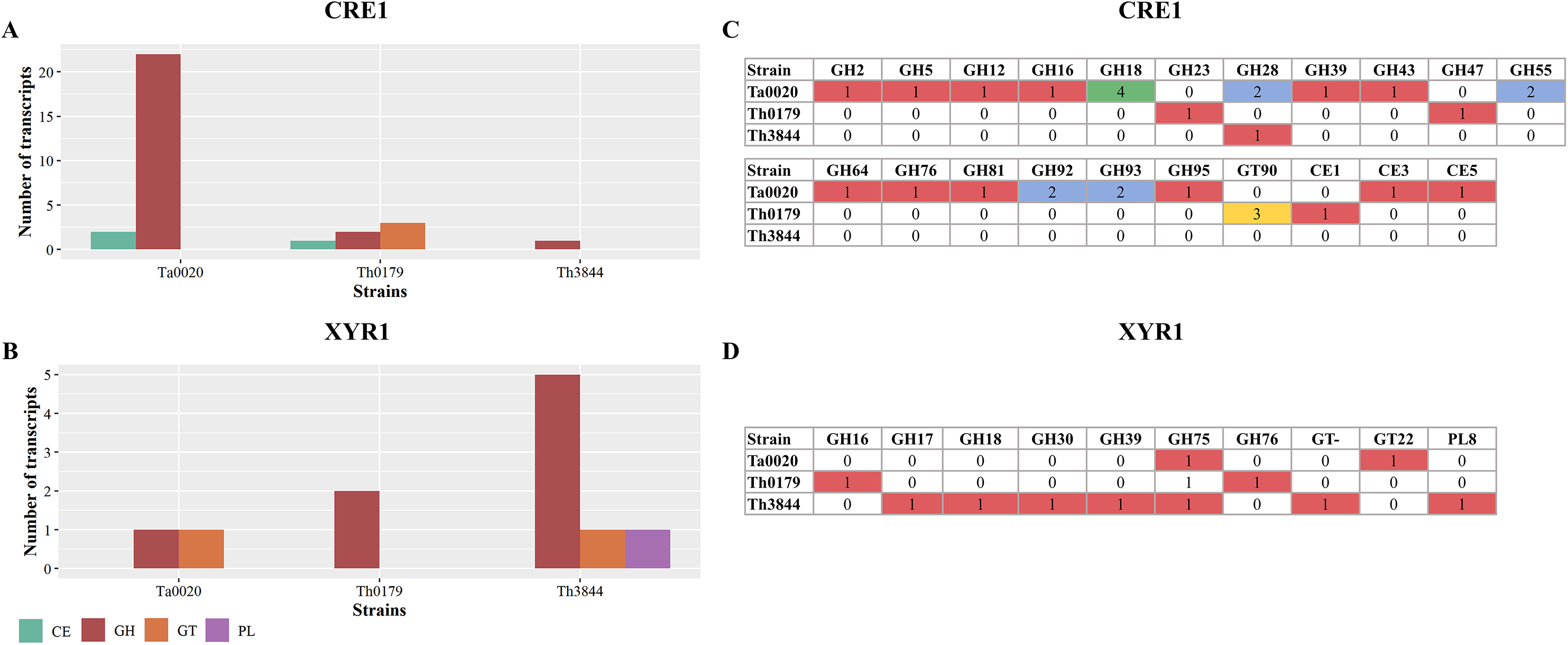
Distribution of CAZyme families in the *cre1* and *xyr1* groups. Classification of CAZyme families coexpressed with *cre1* (**A**) and *xyr1* (**B**) in Th3844, Th0179, and Ta0020 and quantification of each CAZyme family in the *cre1* groups (**C**) and *xyr1* groups (**D**). Illustration showing that the *xyr1* and *cre1* groups presented CAZymes from the GH family in all the evaluated strains with different numbers of transcripts. Overall, Ta0020 had a high number of GHs belonging to family 18, coexpressed with the *cre1* transcript, while Th0179 exhibited a significant amount of GT90. Different CAZyme profiles were observed in the *xyr1* groups, and Th3844 showed the highest diversity of CAZyme families. PL: polysaccharide lyase; GH: glycoside hydrolase; GT: glycosyltransferase; CE: carbohydrate esterase; Ta0020: *T. atroviride* CBMAI-0020; Th0179: *T. harzianum* CBMAI-0179; Th3844: *T. harzianum* IOC-3844.

Several positive and negative regulators are involved in the expression of CAZyme-coding genes and other proteins required for lignocellulose breakdown. Thus, we sought to identify transcripts encoding TFs coexpressed in the *cre1* and *xyr1* groups (Figure 5). In both groups, transcripts encoding Zn_2_Cys_6-_type TFs were identified in all the evaluated strains with different numbers of transcripts. In the *cre1* group of Ta0020, the number was higher than that observed in Th0179 and Th3844 (Figure 5A), while in the *xyr1* group, we observed the opposite profile (Figure 5B). Most Zn_2_Cys_6_ proteins also contain a fungal-specific TF domain, which was coexpressed with both *cre1* and *xyr1* transcripts in Th3844 and Ta0020. In this last strain, a C6 zinc finger domain TF was coexpressed with *xyr1*. Transcripts encoding C_2_H_2-_type TFs were identified in Ta0020 and *T. harzianum* strains in the *cre1* and *xyr1* groups, respectively. Furthermore, in the *xyr1* group, we found a transcript encoding the SteA regulator, which is involved in the regulation of fungal development and pathogenicity (Hoi and Dumas 2010), in Th0179.

**Figure 5.**
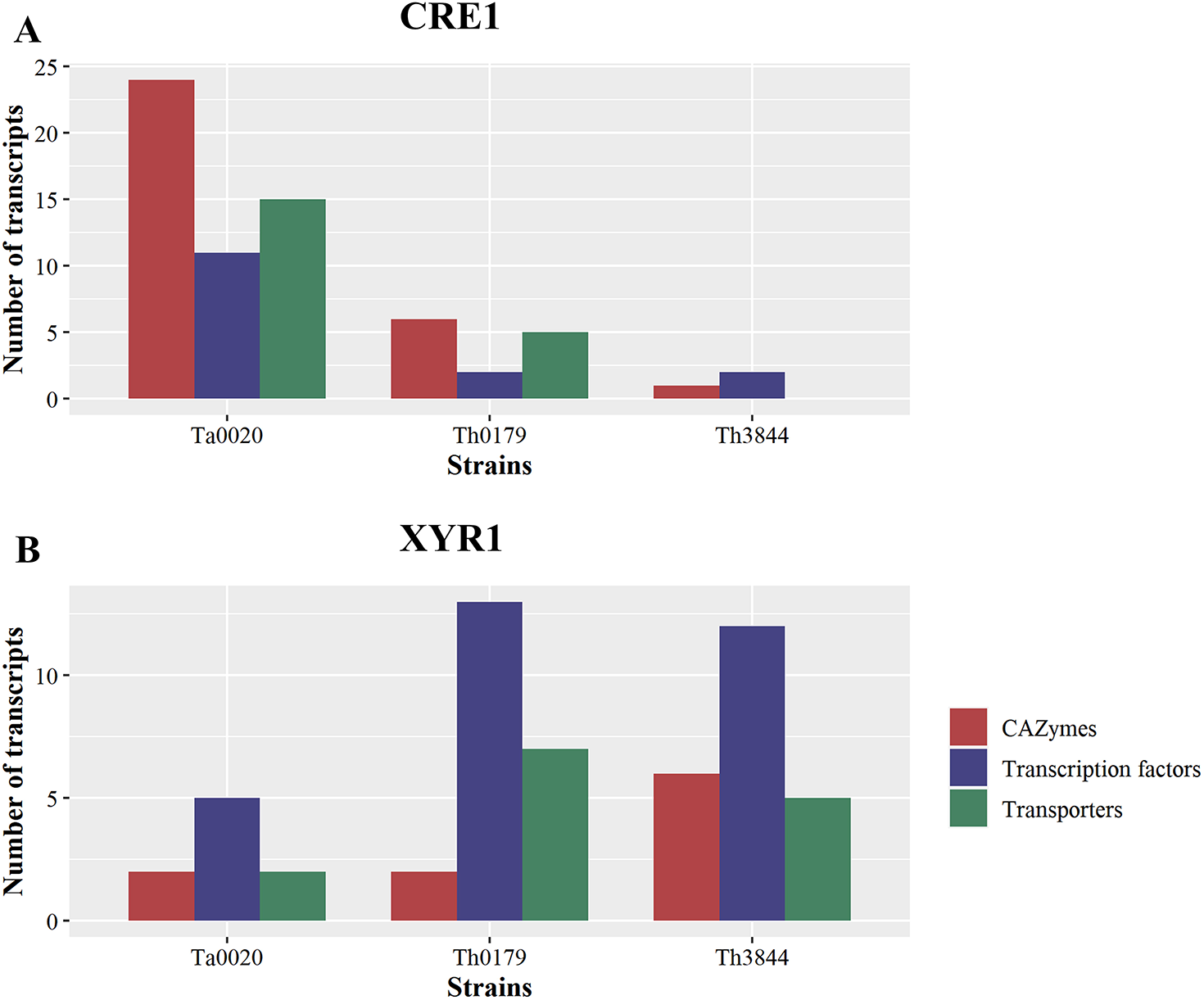
Distribution of TFs and transporters in the *cre1* and *xyr1* groups in *Trichoderma* spp. Classification of TFs coexpressed with *cre1* (**A**) and *xyr1* (**B**) and transporters coexpressed with *cre1* (**C**) and *xyr1* (**D**) in Th3844, Th0179, and Ta0020. The TFs and the transporters were classified according to their class and quantity. MFS: major facilitator superfamily; ABC: ATP-binding cassette; Th3844: *T. harzianum* IOC-3844; Th0179: *T. harzianum* CBMAI-0179; Ta0020: *T. atroviride* CBMAI-0020.

Transporters are also important players in the degradation of plant biomass. Here, transcripts encoding transport proteins coexpressed with *cre1* and *xyr1* were selected (Figure 5). Among them, transcripts encoding major facilitator superfamily (MFS) transport proteins were coexpressed with *cre1* in Th0179 and Ta0020 (Figure 5C) and *xyr1* in Th0179 and Th3844 (Figure 5D). Amino acid transporters were present in the *xyr1* groups of all strains and the *cre1* group only of Th0179. Transcripts encoding the ATP-binding cassette (ABC) were coexpressed with *xyr1* only in Th3844. Th0179 showed transcripts encoding several types of transporters coexpressed with *xyr1*, e.g., vesicle transporter SEC22, cation transporting ATPase, and calcium-dependent mitochondrial carrier, while Ta0020 coexpressed with *cre1*, e.g., UDP-galactose transporter and CNT. Transcripts encoding ion transporters were present in both the *cre1* and *xyr1* groups of Ta0020 and Th0179. In the *cre1* group, such transcripts included zinc and magnesium transporter proteins in Ta0020 and potassium transporter proteins in Th0179. In the *xyr1* group, such transcripts include Cd^2+^Zn^2+^ transporters in Ta0020 and Ca^2+^ transporters in Th0179. Th0179 and Th3844 also present a G protein coexpressed with the *cre1* transcript.

To better understand the regulation mediated by the studied TFs in the evaluated *T. harzianum* strains, we also investigated other classes of proteins coexpressed with the *cre1* and *xyr1* transcripts. For example, phosphatases, kinases and TFs are key components in cellular signaling networks. Transcripts encoding kinase proteins were coexpressed with *cre1* transcripts in Ta0020 and Th3844 and *xyr1* transcripts in Ta0020 and Th0179. In addition, all strains showed transcripts encoding phosphatases coexpressed with *cre1*, and in Th0179 and Th3844, such transcripts were coexpressed with *xyr1*. In addition, transcripts encoding cytochrome P450 genes, which constitute an important group of enzymes involved in xenobiotic degradation and metabolism, were found in the *cre1* groups of all the evaluated strains.

Furthermore, different quantities of differentially expressed transcripts were found in Ta0020 (78) and Th0179 (6) (described in Supplementary Material 6: Supplementary Table 6 and Supplementary Material 1: Supplementary Table 7). In Ta0020, most upregulated transcripts under cellulose growth (70) were associated with the *cre1* group, including *cre1*, whereas *xyr1* demonstrated a potentially significant expression level under glucose growth. In Th3844, *cre1* significantly modulated the expression of the two carbon sources, with a high expression level under glucose growth. Even with the reduced differentiation observed (−1.13), this phenomenon was considered in other studies (Antonieto, et al. 2014; Castro, et al. 2016). In Th0179, *xyr1* and *cre1* revealed a low modulation in transcript expression between growth under cellulose or glucose. However, *xyr1* had a higher transcript expression level under glucose than *cre1*, which had a higher transcript expression level under cellulose growth conditions. In Th3844, *xyr1* exhibited similar results.

### 3.4 Assessing the groups’ network topologies

To identify the main aspects of the network topologies, we applied the HRR methodology to the transcriptome data, which were also used to model the networks using the WGCNA methodology (Figure 6A-C).

**Figure 6.**
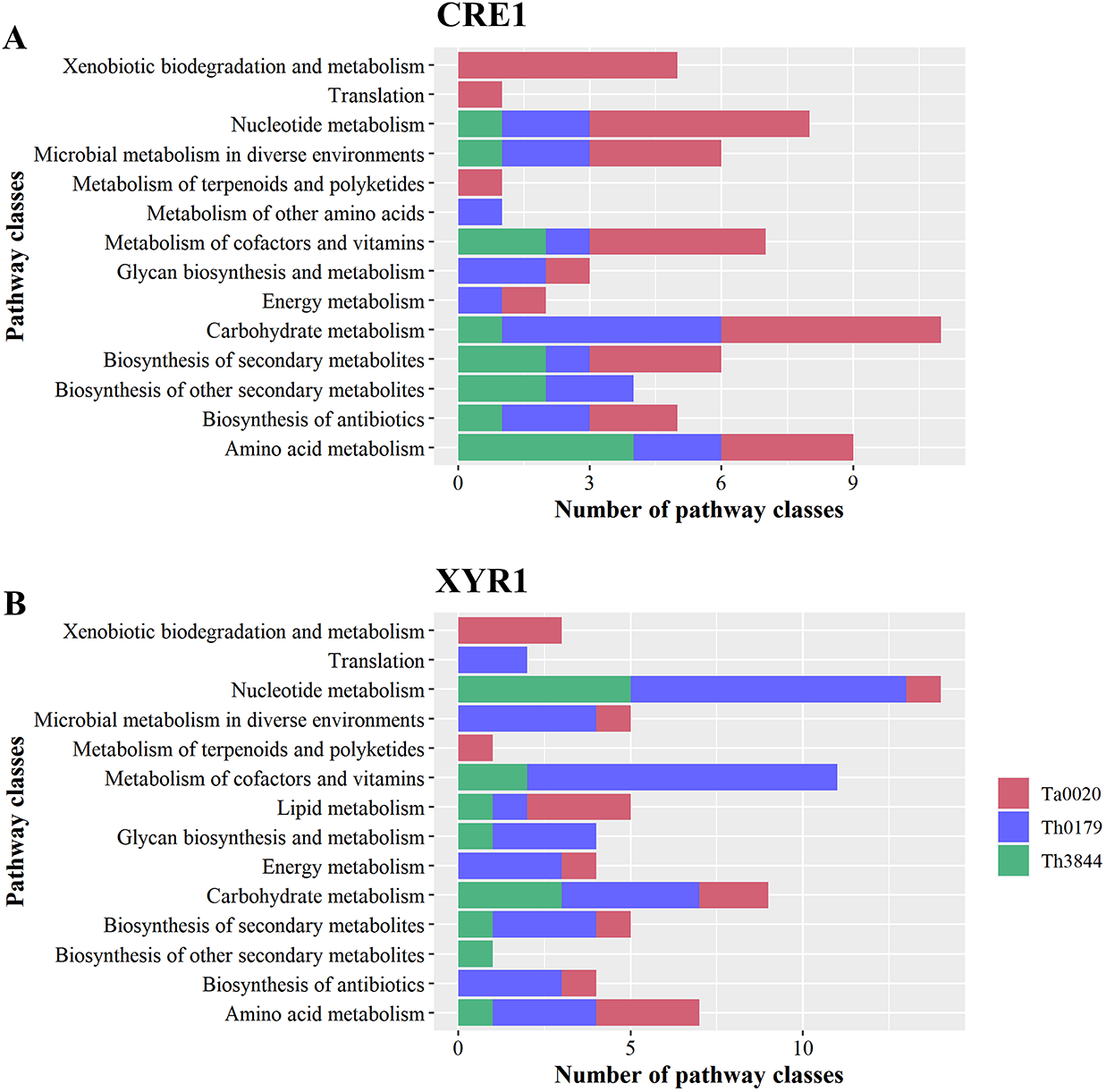
Modeled global networks and modeled subnetworks of the *cre1* and *xyr1* groups of the evaluated strains. The transcriptome datasets were used to infer the global HRR networks of (**A**) Th3844, (**B**) Th0179, and (**C**) Ta0020. The networks were modeled and edited using Cytoscape software. The global networks were partitioned, and subnetworks of Th3844 (**D**), Th0179 (**E**) and Ta0020 (**F**) related to *cre1* were constructed, as well as Th3844 (**G**), Th0179 (**H**) and Ta0020 (**I**) related to *xyr1*. Each color represents a category of transcripts identified in the global networks and subnetworks. Th3844: *T. harzianum* IOC-3844; Th0179: *T. harzianum* CBMAI-0179; Ta0020: *T. atroviride* CBMAI-0020; TF: transcription factor.

**Figure 7.**
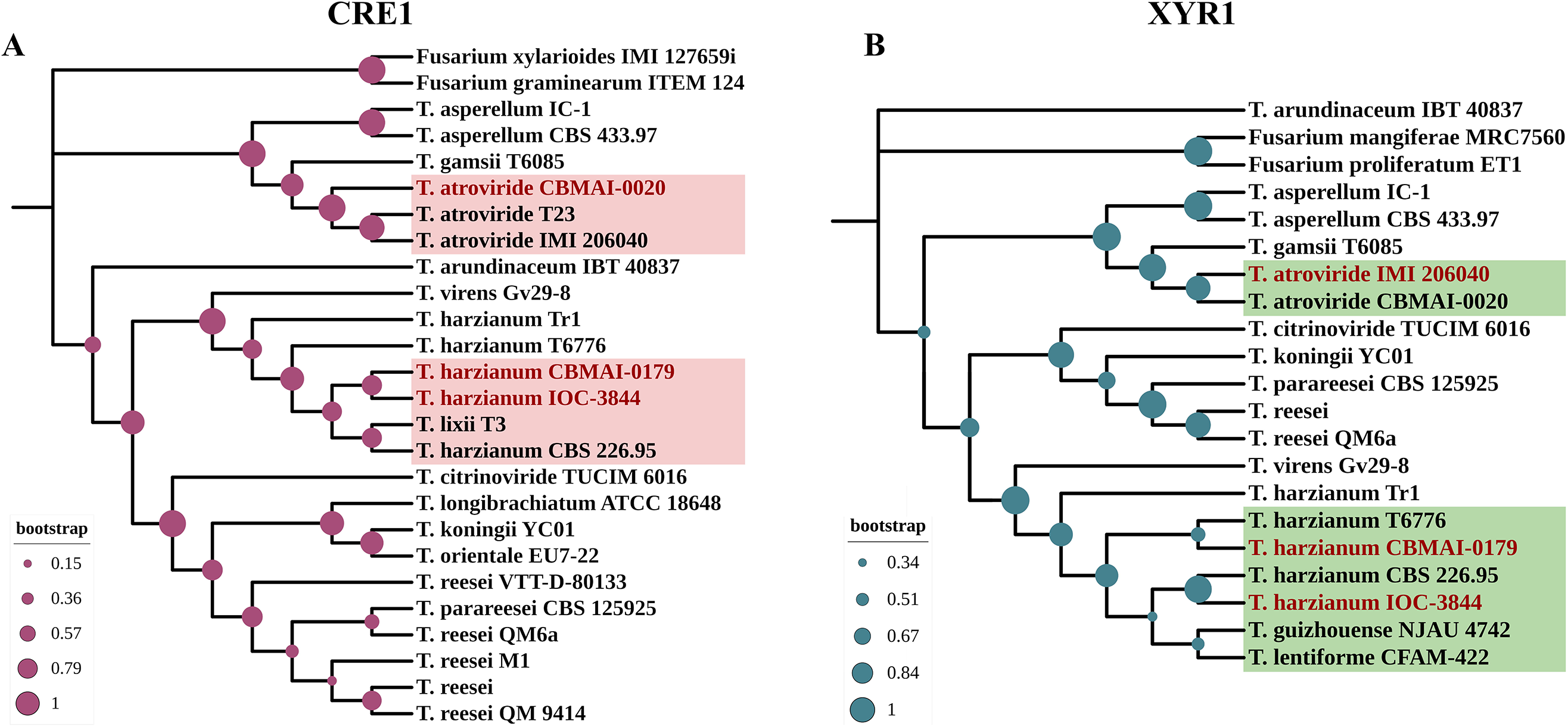
Heatmap plotted based on the shortest pathway between XYR1 or CRE1 and CAZymes in all evaluated *Trichoderma* spp. Number of shortest pathways between XYR1 or CRE1 and the selected CAZymes (**A**) and number of occurrences of such an event (**B**), as indicated by the accentuation of the purple color in the subtitle of the figures. Th3844: *T. harzianum* IOC-3844; Th0179: *T. harzianum* CBMAI-0179; Ta0020: *T. atroviride* CBMAI-0020.

In the CRE1 and XYR1 subnetworks, different connection profiles were observed in each strain (Figure 8). While the transcripts in the *cre1* groups showed less ranked connections in the WGCNA groups of the *T. harzianum* strains Figure 6D and E), Ta0020 showed the opposite profile. In this strain, the transcripts in the *cre1* group were more strongly connected (Figure 6F). In contrast, in the *xyr1* groups, the transcripts were more densely connected in the *T. harzianum* strains (Figure 6G and H), while in Ta0020, the transcripts were separated (Figure 6I).

Transcripts at the top of the degree distribution, which are defined here as hubs, are topologically important to the network structure and are often functionally relevant (Luscombe et al. 2004). Thus, the hub nodes were sought in the *cre1* and *xyr1* groups. In the *cre1* groups, transcripts encoding a SET-domain protein, an ATP synthase, and a transcript not yet annotated were hub nodes of Th3844, Th0179, and Ta0020, respectively. In Ta0020, we found that such an uncharacterized transcript had a degree value equal to 52, which differs from the *T. harzianum* strains in which the hub nodes had degree values of 4 (Th3844) and 9 (Th0179) (Supplementary Material 7: Supplementary Table 8). In the *xyr1* groups, transcripts encoding an ATP-dependent RNA helicase, a cytokinesis sepA, and hypothetical proteins were found as hub nodes of Th3844, Th0179, and Ta0020, respectively (Supplementary Material 7: Supplementary Table 8). Here, we show that the hub nodes in the *cre1* group of Ta0020 had a degree value of 1 and all encoded hypothetical proteins, while in the *T. harzianum* strains, the hub nodes had degree values of 13 (Th3844) and 7 (Th0179). Among the other transcripts with a degree value lower than that of those at the top of the degree distribution, in the *xyr1* group, GH76 and Zn_2_Cys_6-_type TFs (degree value of 6) were identified as candidate hub genes of Th0179, while regulator-nonsense transcripts (degree value of 9) and Zn_2_Cys_6_-type TFs (degree value of 8) were identified as candidate hub genes of Th3844. In the *cre1* group, the proteasome component PRE3 (degree value of 2) in Th3844 was found as a hub node, and replication factor C and phosphoglucomutase (degree value of 5) were found in Th0179. In addition, in Ta0020, we identified calcium calmodulin-dependent kinase (degree value of 9) and GH93 (degree value of 8) as hub nodes.

In all evaluated strains, as a reduced number of differentially expressed transcripts was found in the *cre1* and *xyr1* groups, we expanded our investigation of the studied groups by selecting the first neighbors of XYR1 and CRE1 in global HRR networks. Different quantities of *cre1* neighbors were found in Th3844 (59), Th0179 (21), and Ta0020 (25), and different quantities of *xyr1* were found in Th3844 (46), Th0179 (59), and Ta0020 (70) (described in Supplementary Material 8: Supplementary Table 9). Regarding the *cre1* neighbors, these transcripts were distributed among 24 groups in Th3844, 15 groups in Th0179, and 16 groups in Ta0020. In contrast, considering the *xyr1* neighbors, such transcripts were distributed among 19 groups in Th3844, 32 groups in Th0179, and 42 groups in Ta0020. The direct coexpressed neighbors of *xyr1* included downregulated transcripts in Th3844 (4), Th0179 (5), and Ta0020 (22) and upregulated transcripts in Th3844 (1), Th0179 (1), and Ta0020 (1). Interestingly, only Ta0020 showed differentially expressed transcripts as the first neighbor of the *cre1* transcript (4 downregulated and 4 upregulated). Overall, these differentially expressed transcripts included transporters, CAZymes, kinases, and other proteins related to the regulation process.

For simplification, only a few neighboring transcripts of *xyr1* and *cre1* in Th3844, Th0179, and Ta0020 are shown in Table 1. In Th3844, TFs, transporters, a cytochrome, and a phosphatase were found to be first neighbors of the *cre1* transcript, while TFs, transporters, and CAZymes were found to be the first neighbors of the *xyr1* transcript. In Th0179, several types of proteins, including a kinase, were found to be first neighbors of *cre1*, while transporters, CAZymes, and a TF were found to be first neighbors of the *xyr1* transcript. In Ta0020, several types of proteins, including kinases, were found to be first neighbors of *cre1*, while transporters, CAZymes, TFs, and a DNA ligase were found to be first neighbors of the *xyr1* transcript. Hypothetical proteins were also found to be neighbors of the *cre1* and *xyr1* transcripts of all strains, indicating that both regulators might have a regulatory influence on uncharacterized proteins.

**Table 1.**
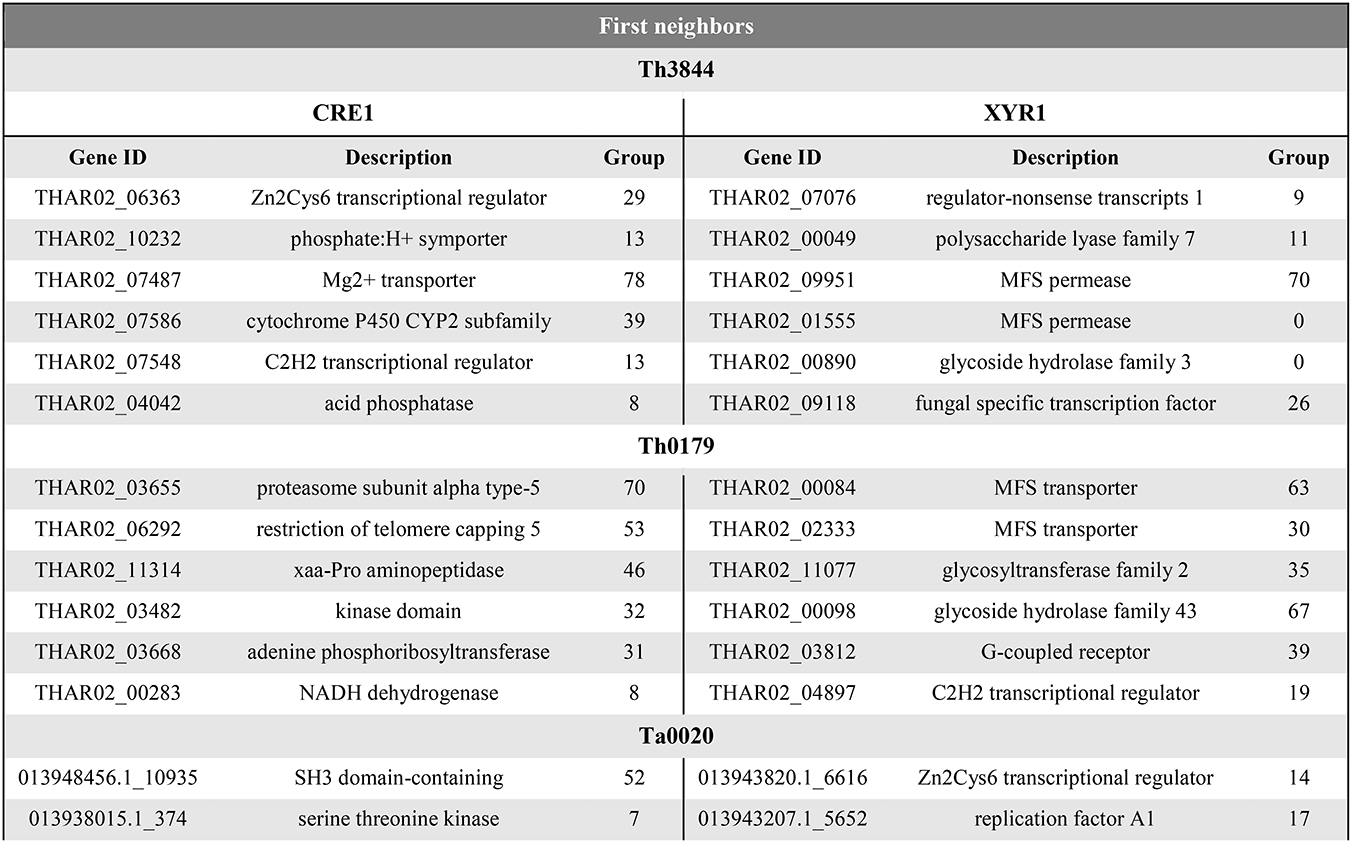
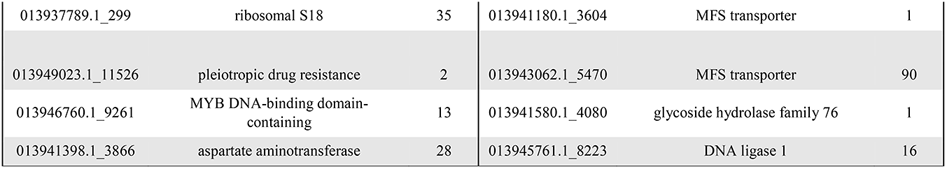
First neighbors of the transcripts *xyr1* and *cre1* identified in Th3844, Th0179, and Ta0020.

| First neighbors |  |  |  |  |  |
| --- | --- | --- | --- | --- | --- |
| Th3844 |  |  |  |  |  |
| CRE1 |  |  | XYR1 |  |  |
| Gene ID | Description | Group | Gene ID | Description | Group |
| THAR02_06363 | Zn2Cys6 transcriptional regulator | 29 | THAR02_07076 | regulator-nonsense transcripts 1 | 9 |
| THAR02_10232 | phosphate:H+ symporter | 13 | THAR02_00049 | polysaccharide lyase family 7 | 11 |
| THAR02_07487 | Mg2+ transporter | 78 | THAR02_09951 | MFS permease | 70 |
| THAR02_07586 | cytochrome P450 CYP2 subfamily | 39 | THAR02_01555 | MFS permease | 0 |
| THAR02_07548 | C2H2 transcriptional regulator | 13 | THAR02_00890 | glycoside hydrolase family 3 | 0 |
| THAR02_04042 | acid phosphatase | 8 | THAR02_09118 | fungal specific transcription factor | 26 |
| Th0179 |  |  |  |  |  |
| THAR02_03655 | proteasome subunit alpha type-5 | 70 | THAR02_00084 | MFS transporter | 63 |
| THAR02_06292 | restriction of telomere capping 5 | 53 | THAR02_02333 | MFS transporter | 30 |
| THAR02_11314 | xaa-Pro aminopeptidase | 46 | THAR02_11077 | glycosyltransferase family 2 | 35 |
| THAR02_03482 | kinase domain | 32 | THAR02_00098 | glycoside hydrolase family 43 | 67 |
| THAR02_03668 | adenine phosphoribosyltransferase | 31 | THAR02_03812 | G-coupled receptor | 39 |
| THAR02_00283 | NADH dehydrogenase | 8 | THAR02_04897 | C2H2 transcriptional regulator | 19 |
| Ta0020 |  |  |  |  |  |
| 013948456.1_10935 | SH3 domain-containing | 52 | 013943820.1_6616 | Zn2Cys6 transcriptional regulator | 14 |
| 013938015.1_374 | serine threonine kinase | 7 | 013943207.1_5652 | replication factor A1 | 17 |
| 013937789.1_299 | ribosomal S18 | 35 | 013941180.1_3604 | MFS transporter | 1 |
| 013949023.1_11526 | pleiotropic drug resistance | 2 | 013943062.1_5470 | MFS transporter | 90 |
| 013946760.1_9261 | MYB DNA-binding domain-containing | 13 | 013941580.1_4080 | glycoside hydrolase family 76 | 1 |
| 013941398.1_3866 | aspartate aminotransferase | 28 | 013945761.1_8223 | DNA ligase 1 | 16 |

We also applied a shortest-path network analysis to identify possible transcripts outside of the *xyr1* and *cre1* groups that may be influenced by CRE1 and XYR1 (described in Supplementary Material 9: Supplementary Table 10). As the shortest path, we considered the minimal number of edges that need to be traversed from a node to reach another node (Koutrouli, et al. 2020). As both regulators act on the gene expression of hydrolytic enzymes related to plant biomass degradation, we sought to determine the number of shortest paths from XYR1 and CRE1 to the selected CAZymes. In all evaluated strains and both TFs, we found a similar quantity of shortest paths (Figure 9A and Supplementary Material 1: Supplementary Figures 6 and 7). However, some aspects should be noted. For example, Th0179 presented transcripts encoding CAZymes from the GH1, GH5, and GH6 families, which are more distantly related to CRE1, while Ta0020 showed only a CAZyme from the GH6 family satisfying this requirement. In Th3844, we found a transcript encoding a CAZyme from GH3 strongly related to XYR1, and CAZymes with the CBM1 domain were also closely connected with such a regulator. In Th0179, a CAZyme from GH55 exhibited a close relationship with XYR1, while in Ta0020, CAZymes from the GH7, GH72, and GH1 families and the CBM1 domain also presented such a profile. In addition, given a determined shortest path, we also considered the number of possibilities that may have occurred (Figure 9B and Supplementary Material 1: Supplementary Figure 6 and 7). Across all strains and in both TFs, significant differences were observed in the number of chances. For instance, a significant number of possible minimum paths between CRE1 and a CAZyme from the GH6 family were observed in Ta0020, likely suggesting a strong influence of this repressor regulator on enzyme activity. Similarly, CAZymes from the AA9 and GH16 families appeared to be affected by XYR1. In Th0179, CAZymes from GH10, GH6, and GH1 presented a higher number of possible shortest paths related to CRE1, potentially indicating an impact of this TF on these enzyme activities.

We also performed a shortest-path analysis to identify transitive transcripts between *cre1* or *xyr1* and the selected CAZymes, which allowed us to discover new transcripts that may be involved in the same biological process. For simplification, we only chose a few shortest paths to explore such transitive transcripts. In Th3844, NADH:ubiquinone oxidoreductase (THAR02_08259), which is the largest multiprotein complex of the mitochondrial respiratory chain (Whitehouse, et al. 2019), was found between GH45 with a CBM1 domain (THAR02_02979) and *cre1*. We also found another transcript involved in ATP synthesis, NAD(P) transhydrogenase beta subunit (THAR02_08260), between GH1 (THAR02_05432) and *cre1*. Furthermore, a transcript encoding a component of the endoplasmic reticulum quality control system called ER-associated degradation (THAR02_08220) (Phillips, et al. 2020) was found between GH16 (THAR02_03302) and *cre1*. Interestingly, in Ta0020, a phosphotyrosine phosphatase was found in the shortest path between GH1 (TRIATDRAFT_135426), which is a homolog of GH1 (THAR02_05432) in *T. harzianum* strains, and *cre1*. This phosphatase is involved in protein phosphorylation, which is a key posttranslational modification critical for the control of many cellular functions (Nasa and Kettenbach 2018). In Th3844, we found a transcript encoding a mitochondrial outer membrane porin (THAR02_07078) between CBM1 (THAR02_02133) and *xyr1*. These protein transporters transport small molecules and play significant roles in diverse cellular processes, including the regulation of mitochondrial ATP and calcium flux (Grevel and Becker 2020). In Ta0020, methyltransferase domain-containing (TRIATDRAFT_292180), which is important for the regulation of chromatin and gene expression (Kouzarides 2007), was found between GH1 (TRIATDRAFT_150220) and *xyr1*.

## 4 Discussion

Although the importance of XYR1 and CRE1 in the expression of CAZyme-encoding genes and other proteins required for lignocellulose degradation is evident, the transcriptional regulation mediated by both proteins in *T. harzianum* strains remains poorly explored (Delabona, et al. 2017, 2021). A previous study demonstrated that some genes encoding regulatory proteins, such as *xyr1*, evolved by vertical gene transfer in *Trichoderma* spp. (Druzhinina, et al. 2018). Here, we inferred the evolutionary relationships of XYR1 and CRE1 in the evaluated strains. In both regulatory proteins, a great genetic distance was observed between the amino acid sequences of Ta0020 and those of Th3844 and Th0179. These findings supported the genetic tree of the evaluated species in which Ta0020 was grouped with other *T. atroviride* strains. However, among the *T. harzianum* strains, we noted that the CRE1 amino acid sequences were more highly conserved than those of XYR1. These results are consistent with previous studies reporting that while the function of CRE1 is conserved throughout the fungal kingdom (Adnan, et al. 2017), the role of XYR1 greatly differs across ascomycete fungi (Klaubauf, et al. 2014).

Recently, different types of enzymatic profiles across *Trichoderma* species were reported, and Th3844 and Th0179, which have hydrolytic potential, have higher cellulase activity during growth on cellulose than Ta0020 (Almeida, et al. 2021). Because such diversity in the enzyme response can be affected by the specific functionalities of regulatory proteins and evolutionary divergence between XYR1 and CRE1 (Benocci, et al. 2017; Klaubauf, et al. 2014), we aimed to investigate how both TFs could affect the activities of the transcripts in response to cellulose degradation in the *T. harzianum* strains. Because transcripts that share the same function or are involved in the same regulatory pathway tend to present similar expression profiles and, hence, form modules (Wolfe, et al. 2005), networks were modeled for Th3844, Th0179, and Ta0020, and the last strain was used to assess the differences across *Trichoderma* species. In fungi, such methodology has been successfully used to provide insight into the regulatory mechanisms of hydrolysis (Arntzen, et al. 2020; Borin, et al. 2018; Li, et al. 2020a).

In recent years, network methodologies have been employed for the discovery of new targets associated with biological processes (Cortijo, et al. 2020; Liu, et al. 2021; Zhang, et al. 2021). Because such approaches might provide mechanisms for inferring new gene relationships and functions, experimental validation by the construction of mutants is usually a way of corroborating these insights. In this context, the construction of mutants can show the influence of a unique missing gene on genome regulation, helping to corroborate the hypotheses raised in the present study. However, due to the inefficient homologous recombination machinery of filamentous fungi, once a fungus reproduces asexually, a low frequency of correct genomic integration has been observed for *Trichoderma* spp. (Liu, et al. 2017; Vieira, et al. 2018; Zeilinger 2004). These challenges can be overcome by using the CRISPR/Cas9 system, which represents a good alternative in *Trichoderma* spp. (Fonseca, et al. 2020; Vieira, et al. 2021) and may enable the future establishment of efficient protocols for each strain described here. Thus, by predicting the profiles of the transcripts coexpressed with XYR1 and CRE1, the results can be used to engineer mutant strains with the objective to investigate the influence of the regulation of such TFs *in vitro*.

We observed that each network had a different configuration with distinct module profiles of XYR1 and CRE1, providing insight into how these TFs might act in specific ways in different *Trichoderma* species. These functional differences in the *xyr1* and *cre1* groups can be attributed to previously reported differences in the regulatory mechanisms of hydrolysis (Almeida, et al. 2021). By analyzing both groups, we observed that more transcripts were coexpressed with XYR1 in the *T. harzianum* strains than *T. atroviride*, which had more transcripts coexpressed with CRE1. However, although the phylogenetic similarity between the evaluated *T. harzianum* strains shows that they share a largely common genetic background, their profiles of transcripts coexpressed with *xyr1* and *cre1* differed.

### 4.1 Insight into the genetic impacts of XYR1 and CRE1

To obtain insight into the functional profile of the *cre1* and *xyr1* groups, GO enrichment analyses of transcripts from these groups of all evaluated strains were performed. Here, response to external stimulus was a notable GO term in Th0179 in the *xyr1* group, suggesting that the transcripts of these group likely respond to external environmental conditions, such as carbon sources. Furthermore, interconnections between nutrient and light signaling pathways have been reported in filamentous fungi, such as *N. crassa* and *T. reesei*, with substantial regulation by photoreceptors (Schmoll 2018). More interestingly, the influence of light on CRE1 functions has been reported (Monroy, et al. 2017), supporting our findings showing that the response to a light stimulus was an enriched GO term in the *cre1* group of Th3844. In contrast, the enrichment analysis of Ta0020 indicates that the activity of CRE1 may be stronger in such fungi, directly repressing genes related to plant cell wall-degrading enzymes, which was not observed in the *T. harzianum* strains. In these strains, organic substance metabolic processes (Th3844) and regulation processes (Th0179) were enriched GO terms. In addition, fungal-type cell wall organization was an enriched term of Th0179 in the *cre1* group.

By investigating the KO functional annotation of the *cre1* and *xyr1* groups, we suggest that both regulators act on the same enzymatic pathways, which was expected due to antagonism in their function, i.e., while CRE1 is the main repressor of genes encoding proteins related to lignocellulose degradation, XYR1 is the main activator of such genes. However, each triggers a specific metabolic pathway according to the strain; therefore, some aspects are noteworthy. For example, it has been reported that the *Trichoderma* species display several mechanisms during their antagonistic action against plant pathogens, including the production of secondary metabolites (Malmierca, et al. 2015). According to our findings, due to the highest number of pathways related to secondary metabolites, including the metabolism of terpenoids and polyketides (Mukherjee, et al. 2008), we suggest that CRE1 and XYR1 participate in the regulation of secondary metabolism compounds in *T. atroviride*. Furthermore, XYR1 in *T. harzianum* strains appeared to be deeply connected with enzyme pathways related to the metabolism of carbohydrates, especially in the Th3844 strain.

Through a network analysis, we identified transcripts encoding CAZymes coexpressed with *xyr1* and *cre1* in all evaluated strains. We identified CAZyme families responsible for cellulose degradation (e.g., GH12); hemicellulose degradation (e.g., GH16, GH17, CE1, and CE5); pectin degradation (e.g., GH28 and GH93); and other CAZy families with multiple activities or minor activities on lignocellulosic substrates, such as GH2, GH5, GH43, GH95, GH30, and GH39 (de Vries, et al. 2017; Kameshwar, et al. 2019). Although Ta0020 presented the highest number of GHs coexpressed with CRE1, it was found to be less efficient than the other *Trichoderma* strains in degrading plant biomass (Almeida, et al. 2021). In the GH class, enzymes belonging to the GH18 family, which is mainly represented by chitinase-like proteins, are directly related to fungal cell wall degradation in mycoparasite species of *Trichoderma* (Gruber and Seidl-Seiboth 2012). Four GH18 enzymes were coexpressed with CRE1 in *T. atroviride*, which is widely used as a biocontrol agent in agriculture. However, in Ta0020, enzymes from the GH75 family of chitosanases were coexpressed with *xyr1*. Therefore, the degradation of chitosan, which is a partially deacetylated derivative of chitin (Hahn, et al. 2020), is a relevant aspect of mycoparasitism that may be influenced by the activity of XYR1. Recently, transcripts encoding GHs were found to be coexpressed with some TFs involved in biomass degradation, e.g., XYR1 (Borin, et al. 2018). In the present study, the transcripts encoding hydrolytic enzymes could be sorted into other groups formed in the networks mainly because their expression patterns differed from those of the *xyr1* and *cre1* transcripts.

Several fungal TFs have been described to be directly involved in the regulation of plant biomass utilization (Benocci, et al. 2017). Most TFs belong to the zinc cluster family, including Zn_2_Cys_6-_ and C_2_H_2_-type TFs, which are characterized by the presence of zinc finger(s) in their binding domains. Most positive regulators appear to belong to the Zn_2_Cys_6_ class, while repressors belong to the C_2_H_2_ class (Benocci, et al. 2017). Here, we found the highest number of transcripts encoding the Zn_2_Cys_6_ and C_2_H_2_ classes in the *xyr1* and *cre1* groups of Th0179 and Ta0020, respectively. Both *T. harzianum* strains had a similar profile of transcripts coexpressed with CRE1 and XYR1. However, SteA, a C_2_H_2-_type TF, was coexpressed with the *xyr1* transcript of Th0179. This regulator has been described as an important player in fungal environmental adaptation in response to nutrient deprivation, the production of extracellular proteins involved in the degradation of complex substrates (Hoi and Dumas 2010), and the mediation of the regulatory role of mitogen-activated protein kinase (MAPK) during mycoparasitic responses (Gruber and Zeilinger 2014). In *Aspergillus nidulans*, SteA is required for sexual development (Vallim, et al. 2000).

During plant biomass degradation, fungi secrete extracellular enzymes to decompose polysaccharides into small molecules, which are then imported into cells through transporters (Sloothaak, et al. 2016). One of the most relevant sugar transporter families in filamentous fungi is the MFS family (Zhang, et al. 2013). Here, we found the highest number of transcripts encoding MFS coexpressed with *cre1* and *xyr1* in Ta0020 and Th3844, respectively. Among the *T. harzianum* strains, compared with Th3844, Th0179 showed transcripts encoding several types of transporter proteins, such as transporter proteins of calcium ions (THAR02_04202, THAR02_03989, and THAR02_01350). It has already been reported that metal ions, such as Ca^2+^, have a positive effect on the mycelial growth of *T. reesei* and cellulase production (Chen, et al. 2016). This molecular signaling mechanism is mediated by cations transporting ATPase and calcium-dependent mitochondrial carriers, which are both components of Ca^2+/^calmodulin signal transduction, including the TF Crz1 (Chen, et al. 2016; Martins-Santana, et al. 2020). Therefore, investigating the role of proteins related to calcium transporters in the induction of genes responsive to lignocellulose degradation in Th0179 cells is important since these proteins could transport cations that activate gene expression. Furthermore, transcripts encoding G proteins were coexpressed with *cre1* in Th0179 and Ta0020. Heterotrimeric G proteins have been well studied in several *Trichoderma* species. In saprophytic species, such proteins are involved in the nutrient signaling pathway in connection with a light response, triggering the posttranscriptional regulation of cellulase expression (Hinterdobler, et al. 2021); in mycoparasitic species, G protein-coupled receptors are involved in the regulation of processes related to mycoparasitism (Zeilinger and Atanasova 2020).

Although CAZymes, TFs, and transporters play an important role in cellulose degradation, these types of proteins represented only a small percent of the transcripts coexpressed with XYR1 and CRE1 compared to the other protein classes as follows: in the *cre1* group, (I) 10.4% (Th0179), (II) 5.7% (Th3844), and (III) 17.9% (Ta0020), and in the *xyr1* group, (IV) 11.8% (Th0179), (V) 11.9% (Th3844), and (VI) 16% (Ta0020). Since the regulation of genes involved in biomass breakdown is a complex process that involves several signaling pathways, we expected to find proteins with a great range of functions coexpressed with *xyr1* and *cre1*. For example, various kinases and phosphatases were coexpressed with *xyr1* and *cre1* in all evaluated strains. It has been reported that cellulase gene expression can be regulated by the dynamics of protein phosphorylation and dephosphorylation, which involve protein kinases and phosphatases, respectively (Schmoll, et al. 2016). Furthermore, in filamentous fungi, phosphorylation is a prerequisite for CRE1 activity (Cziferszky, et al. 2002; de Assis, et al. 2021; Han, et al. 2020). Therefore, in *T. harzianum*, it is important to elucidate the role of kinases and phosphatases in the regulation of CRE1 function. In all evaluated strains, cytochrome P450 coding genes represented another class of proteins coexpressed in the *cre1* group. It has been reported that these enzymes are important for cells to perform a wide variety of functions, including primary and secondary metabolism, xenobiotic degradation, and cellular defense against plant pathogenic fungi (Chadha, et al. 2018; Fan, et al. 2013; Siewers, et al. 2005). To expand previous findings concerning *Trichoderma* spp. (Chadha, et al. 2018), their similar expression pattern with the *cre1* transcript makes them important candidates for an extensive investigation in the cellulose degradation context.

We also investigated the expression profiles of the transcripts in the *cre1* and *xyr1* groups, including *xyr1* and *cre1*, under cellulose and glucose growth. Castro, et al. (2016) reported that in *T. reesei*, the expression level of *xyr1* was minimal in the presence of glucose; this phenomenon was not observed in Ta0020 in the present study. In addition, in Ta0020, *cre1* was upregulated in the presence of cellulose, which was not expected due to the role of CRE1 in the repression of cellulolytic and hemicellulolytic enzymes when an easily metabolizable sugar, i.e., glucose, is available in the environment (Benocci, et al. 2017). In contrast, in Th3844, the *cre1* expression level in the presence of glucose was higher than that in the presence of cellulose. The gene coexpression networks were modeled based on the transcript expression level under two sets of conditions (cellulose and glucose). Considering that the expression of the *cre1* transcript in Ta0020 was upregulated under cellulose growth, upregulated transcripts were expected and found in the *cre1* group. Recently, Almeida et al. (Almeida, et al. 2021) reported that Ta0020 presented the highest number of differentially expressed transcripts under cellulose growth conditions relative to glucose, followed by Th3844 and Th0179. Here, the quantity of upregulated transcripts was low in the *cre1* and *xyr1* groups of the *T. harzianum* strains. These results may indicate that both TFs have a basal expression level at 96 h of fermentation; thus, the transcripts grouped with such regulatory proteins presented the same expression pattern and, mostly, were not differentially expressed transcripts.

### 4.2 Examining the network’s topology

To extract additional information from the networks, we characterized the network topologies of the modules identified separately. We might infer that many transcripts were under the influence of the repressor CRE1 in Ta0020, while in the *T. harzianum* strains, the transcripts seem to have been affected by the activator XYR1. Therefore, the described profiles may indicate that in *T. harzianum*, the transcripts, including those encoding XYR1, act together to perform a determined biological function that is favorable to the expression of genes related to cellulose degradation. In contrast, in *T. atroviride*, the transcripts in the *cre1* group appeared to act on the same biological process, which may be related to the repression of genes encoding hydrolytic enzymes and other proteins required for cellulose degradation.

Hub transcripts were identified in the *cre1* and *xyr1* groups, and new targets were discovered. In the *cre1* group, a transcript with a SET-domain coding gene was found as a hub node of Th3844. Such SET-domain proteins participate in chromatin modifications by methylating specific lysines on the histone tails (Kouzarides 2007). In filamentous fungi, the epigenetic regulation of holocellulase gene expression has already been reported (Zeilinger, et al. 2003), and CRE1 plays an important role in nucleosome positioning (Ries, et al. 2014). Interestingly, ATP synthase, which synthesizes ATP from ADP and inorganic phosphate on mitochondria (Burger, et al. 2003), was found to be a hub node of Th0179. Thus, we might infer that a great number of transcripts were coexpressed with a hub node related to mitochondrial ATP production, which is the main energy source for intracellular metabolic pathways (Neupane, et al. 2019). In the *xyr1* group, a transcript encoding an ATP-dependent RNA helicase was found as a hub node of Th3844. Such enzymes catalyze the ATP-dependent separation of double-stranded RNA and participate in nearly all aspects of RNA metabolism (Jankowsky 2011). Additionally, the sepA transcript was identified as a hub node of Th0179. In *Aspergillus nidulans*, the sepA gene encodes a member of the FH1/2 protein family, which is involved in cytokinesis and the maintenance of cellular polarity, i.e., related to the cell division process (Harris, et al. 1997). Overall, our findings suggest that the hub genes were not necessarily the most effective genes related to lignocellulose deconstruction, confirming the indirect action of XYR1 and CRE1 in *T. harzianum* on regulatory hydrolysis mechanisms. In Ta0020, transcripts encoding hypothetical proteins were identified as potential hub nodes in both groups, providing new targets for further studies evaluating their functions in fungal physiology.

We also investigated the first neighbors of the *cre1* and *xyr1* transcripts in the modeled global HRR networks of all evaluated strains. While Ta0020 showed differentially expressed transcripts as the first neighbor of the *cre1* transcript, interestingly, both *T. harzianum* strains presented the opposite profile in which only the first neighbors of the *xyr1* transcript were differentially expressed. In addition, the evaluated strains showed different numbers of neighbors of the *cre1* and *xyr1* transcripts, which were distributed among several groups. Such a divergent profile may indicate that each TF affects a set of transcripts in a specific way that varies among the strains. Overall, we identified CAZymes, TFs, and transporters as first neighbors of *cre1* and *xyr1*, confirming the results obtained in this study using the WGCNA approach.

Another property of a network’s topology is the shortest paths connecting two transcripts (Pavlopoulos, et al. 2011). Here, the significant shortest paths between both studied TFs, i.e., CRE1 and XYR1, and the CAZymes with a higher-level expression under cellulose growth conditions were investigated in all strains. The average shortest pathway distance between the transcripts encoding CRE1 or XYR1 and all selected CAZymes was relatively short, which may be attributed to a network phenomenon called the small-world effect, i.e., networks can be highly clustered with a small number of necessary steps to reach one node from another (Maier 2019). However, several aspects should be highlighted. We identified CAZymes responsible for cellulose degradation, such as GH6 (Th0179 and Ta0020), and other CAZy families with multiple activities or minor activities on lignocellulosic substrates, such as GH1 and GH5 (Th0179) (de Vries, et al. 2017; Kameshwar, et al. 2019) with a low number of minimum pathways to CRE1 in Ta0020. In contrast, CAZymes responsible for cellulose degradation, such as CBM1 domain (Th3844 and Ta0020) and GH7 (Ta0020), for hemicellulose degradation, such as GH55 (Th0179), and CAZy families with multiple activities or minor activities on lignocellulosic substrates, such as GH3 (Th3844) and GH1 (Ta0020) (de Vries, et al. 2017; Kameshwar, et al. 2019), were identified with a high number of minimum pathways to XYR1. Because certain shortest paths are not necessarily unique, we also considered the number of possibilities that the paths may occur. Our results suggest that the number of possible minimum paths between some CAZymes and CRE1 or XYR1 was higher than of others. For example, a great number of possible shortest paths was observed between CRE1 and GH6 in Ta0020 and between CRE1 and GH10, GH6, or GH1 in Th0179. In contrast, in Ta0020, such profiles were observed between XYR1 and CAZymes with cellulolytic (AA9) and hemicellulolytic activities (GH16) and between CRE1 and GH6 with a CBM1 domain.

Furthermore, we performed a shortest path analysis to explore transitive transcripts between two nodes. Considering that the lowest number of transitive transcripts between two nodes may indicate the need for fewer signal pathways, representing a more direct relation, we chose to investigate the shortest paths with only one transcript between the desired targets. Interestingly, transcripts encoding proteins related to ATP synthesis were found between GH45 with a CBM1 domain and *cre1* and between GH1 and *cre1* in Th3844, which could be explained by the demand for energy required for CRE1 to exercise its repressor activity on these hydrolytic enzymes. However, in Th3844, a transcript encoding a protein involved in quality control processes that center on the endoplasmic reticulum was found between GH16 and *cre1*, indicating that a signaling pathway is triggered by CRE1 to repress the expression of such an enzyme. Curiously, in Ta0020, a phosphotyrosine phosphatase was found in the shortest path between GH1, which is a homolog of GH1 in the *T. harzianum* strains, and *cre1*. Such proteins are involved in posttranslational modification, including the phosphorylation process, which plays an essential role in signal transduction to achieve CCR by CRE1 (Han, et al. 2020; Horta, et al. 2019). In Th3844, we found a transcript encoding a protein related to the transport of substances across the mitochondrial membrane between CBM1 and *xyr1*, which may indicate high cellular activity and, therefore, a high demand for energy for gene expression. In Ta0020, a transcript encoding a protein involved in the regulation of chromatin and gene expression was found between GH1 and *xyr1*, which may indicate an intermediated process for the expression of such an enzyme.

In conclusion, biological networks represent a powerful approach to accelerate the elucidation of the molecular mechanisms underlying important biological processes. Our findings suggest that the set of transcripts related to XYR1 and CRE1 varies among the studied *T. harzianum* strains, suggesting regulatory differences in enzymatic hydrolysis. These findings corroborate previous studies in which differences in biomass degradation and enzyme production between strains of the same species have been reported (de Vries, et al. 2017; Thanh, et al. 2019; Tolgo, et al. 2021). Furthermore, such transcripts were not limited to CAZymes and other proteins related to biomass degradation. Thus, we suggest that both TFs play a role in the undirected regulation of gene encoding proteins related to cellulose degradation, and multiple pathways related to gene regulation, protein expression, and posttranslational modifications may be triggered by the studied TFs. We expect that our results could contribute to a better understanding of fungal biodiversity, especially regarding the transcription regulation involved in hydrolytic enzyme expression in *T. harzianum*. By describing new potential targets involved in the cellulose degradation pathway, this knowledge can be used to develop genetic manipulation strategies and expand the use of *T. harzianum* as an enzyme producer in biotechnological industrial applications.

## Author contributions

**RRR**: Writing - original draft, Methodology, and Conceptualization. **AHA**: Methodology, Software, Writing - review & editing, and Formal analysis. **DAA**: Resources, and Writing - review & editing. **JAFF**: Writing - review & editing. **MACH**: Resources and Writing - review & editing. **APS**: Supervision, review & editing, and funding acquisition.

## Abbreviations

ABC: ATP-binding cassette
CAZymes: carbohydrate-active enzymes
CCR: carbon catabolite repression
CEs: carbohydrate esterases
CRE1: carbon catabolite repressor 1
GHs: glycoside hydrolases
GO: Gene Ontology
GTs: glycosyltransferases
HRR: highest reciprocal rank
iTOL: Interactive Tree of Life
ITS: internal transcribed spacer
JTT: Jones-Taylor-Thornton
K2P: Kimura two-parameter
KEGG: Kyoto Encyclopedia of Genes and Genomes
KO: KEGG Orthology
MEGA: Molecular Evolutionary Genetics Analysis
MFS: major facilitator superfamily
ML: Maximum likelihood
PLs: polysaccharide lyases
Ta0020: *Trichoderma atroviride* CBMAI-0020
*tef1*: translational elongation factor 1
TFs: transcription factors
Th0179: *Trichoderma harzianum* CBMAI-0179
Th3844: *Trichoderma harzianum* IOC-3844
TOM: topological overlap matrix
TPM: transcripts per million
WGCNA: weighted correlation network analysis
XYR1: xylanase regulator 1

## Acknowledgments

We are grateful to the Brazilian Biorenewables National Laboratory (LNBR), Campinas – SP for conducting the fermentation experiments; the Center of Molecular Biology and Genetic Engineering (CBMEG) at the University of Campinas, SP for the use of the center and laboratory space; and the São Paulo Research Foundation (FAPESP), the Coordination of Improvement of Higher Education Personnel (CAPES, Computational Biology Program), and the Brazilian National Council for Technological and Scientific Development (CNPq) for supporting the project and researchers.

## Data availability statement

All data generated or analyzed in this study are included in this published article (and its supplementary information files). The raw RNA-Seq datasets were deposited at the NCBI Sequence Read Archive and can be accessed under the BioProject number PRJNA336221.

