## Supplementary Material 1 for "Network Analysis Reveals Different Cellulose Degradation Strategies across *Trichoderma harzianum* Strains Associated with XYR1 and CRE1"

**Rafaela Rossi Rosolen[1,2], Alexandre Hild Aono[1,2], Déborah Aires Almeida[1,2], Jaire Alves Ferreira Filho[1,2], Maria Augusta Crivelente Horta[1], Anete Pereira de Souza[1,3*]**

[1]Center for Molecular Biology and Genetic Engineering (CBMEG), University of Campinas (UNICAMP), Cidade Universitária Zeferino Vaz, Campinas, SP, Brazil

[2]Graduate Program in Genetics and Molecular Biology, Institute of Biology, UNICAMP, Campinas, SP, Brazil

[3]Department of Plant Biology, Institute of Biology, UNICAMP, Cidade Universitária Zeferino Vaz, Rua Monteiro Lobato, 255, Campinas, SP, Brazil

*** Correspondence:**
Anete Pereira de Souza


**Supplementary data**

This PDF file includes the following:

**Supplementary Material 1**

#### Materials and methods

#### Fungal strains, culture conditions, and transcription profiling

#### Results


- **Supplementary Table 1**
- **Supplementary Figure 1**
- **Supplementary Figure 2**
- **Supplementary Figure 3**
- **Supplementary Figure 4**
- **Supplementary Figure 5**
- **Supplementary Figure 6**
- **Supplementary Figure 7**
- **Supplementary Table 7**

### 1 Material and methods

#### 1.1 Culture conditions and transcription profiling

Briefly, the *T. harzianum* CBMAI-0179 (Th0179), *T. harzianum* IOC-3844 (Th3844), and *T. atroviride* CBMAI-0020 (Ta0020) strains were cultivated on solid medium for 8 days at 37 °C to produce a sufficient number of spores as an inoculum for fermentation as described in a previous study (Horta et al., 2018). The fermentation process was initiated with the inoculation of $10^7$ spores/mL in an initial volume of 200 mL of Mandels Andreotti (MA) minimal medium (Mandels and Andreotti, 1978) with crystalline cellulose (Celuflok, São Paulo, Brazil, degree of crystallinity 0.72 g/g, composition 0.857 g/g cellulose and 0.146 g/g hemicellulose) or glucose (control condition) as the carbon source (Horta et al., 2018). After 72 h of incubation, 50 mL of the preinoculum were used to inoculate 450 mL of fermentation solution in 2 L Erlenmeyer flasks in which the composition varied according to the carbon source (Horta et al., 2018). The fermentation process of all evaluated strains was performed in biological triplicates and continued for 96 h. Then, mycelial samples were extracted, stored at -80 °C and ground in liquid nitrogen. Frozen material was used for the RNA extraction, and the sequencing experiment was carried out on an Illumina HiSeq 2500 platform (Illumina, San Diego, CA, USA).

The transcriptome analysis was performed in a previous study conducted by our research group (Almeida et al., 2021). Briefly, the reads obtained by Horta et al. (2018) (BioProject accession PRJNA336221) were size-filtered (minimum length, 36 bp) and selected by quality (Q > 15) using Trimmomatic v0.36 (Bolger et al., 2014). The transcript identification was obtained from comparative alignments of sequencing data along with the *T. harzianum* T6776 (Baroncelli et al., 2015) genome of Th3844 and Th0179 and the *T. atroviride* IMI206040 (Kubicek et al., 2011) genome of Ta0020 as reference genomes using CLC Genomics Workbench software (CLC bio – 6.5.2 v; Finlandsgade, Dk) (CLC, 2016).

To evaluate differences from the reference genome, the reads from the sequencing data were mapped against other public reference genomes of *T. harzianum* strains available in the NCBI GenBank database (*T. harzianum* TR274 (Steindorff et al., 2014) and *T. harzianum* CBS 226.95 (PRJNA207867)). No significant differences were observed between the treatments, supporting the use of the *T. harzianum* T6776 genome as a reference (Santos et al., 2016; Filho et al., 2017; Horta et al., 2018). Furthermore, close phylogenetic proximity was observed among the T6776, CBS 226.95, and TR274 strains of *T. harzianum*, which exhibit similar genome sizes and similar numbers of orthologs and paralogous genes (Kubicek et al., 2019) **(**Supplementary Table 1). The gene expression values were calculated in transcripts per million (TPM). To statistically analyze the differentially expressed genes (DEGs), fold change ≥ 1.5 or ≤ -1.5 and a p-value ≤ 0.05 were applied as previously described (Almeida et al., 2021).

**Supplementary Table 1.** Value (%) of reads mapped against the reference genomes.

| | Reference genome | | |
|---|---|---|---|
| **Strain** | **ThT6776** | **ThTR274** | **ThCBS226.95** |
| **Th0179** | 53.88 | 53.31 | 54.39 |
| **Th3844** | 66.15 | 67.61 | 67.94 |

The value (%) of the sequence mapping was estimated by the alignment of the reads from Th0179 and Th3844 against the reference genomes. Th0179: *T. harzianum* CBMAI-0179; Th3844: *T. harzianum* IOC-3844; ThT6776: *T. harzianum* T6776; ThTR274: *T. harzianum* TR274; and ThCBS226.95: *T. harzianum* CBS 226.95.

#### 2 Results

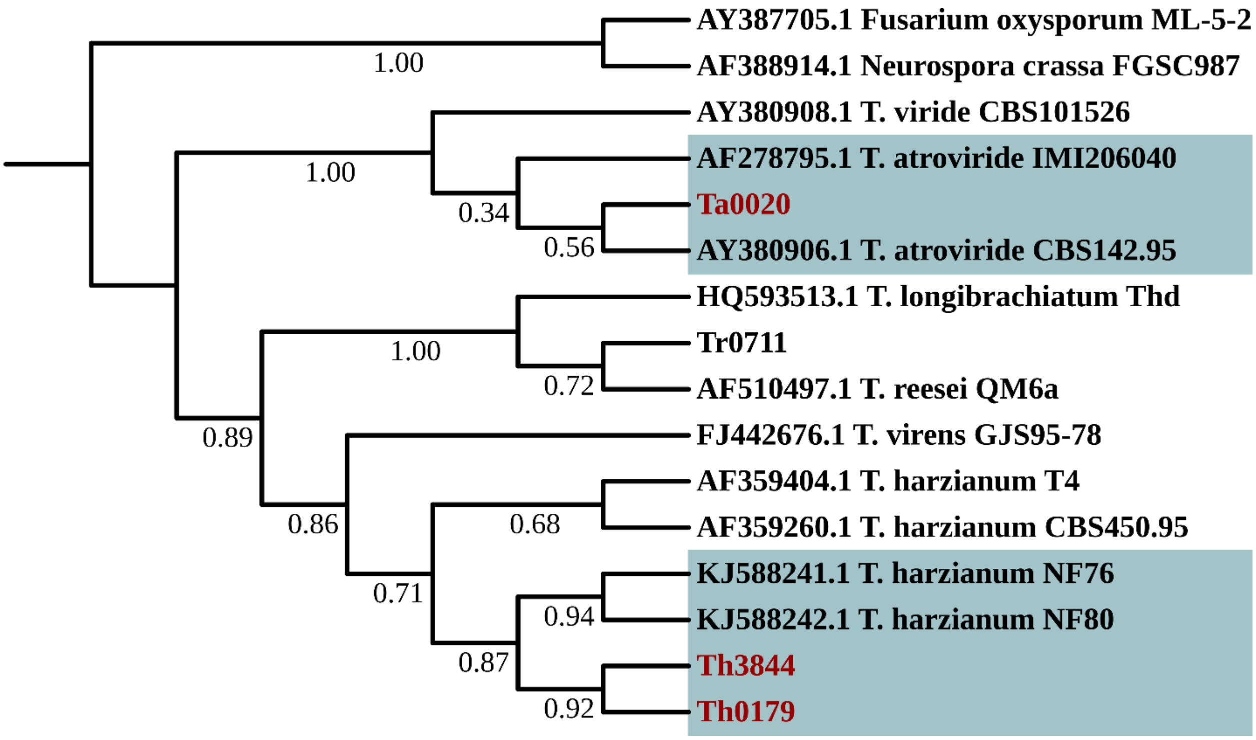


**Supplementary Material 1: Supplementary Figure 1. Phylogenetic relationships of *Trichoderma* spp. inferred by an analysis of the ITS sequences.** Sequences corresponding to the ITS region were used to analyze and compare the phylogenetic relationships of the studied *Trichoderma* species/strains. The ITS sequences of Th3844, Th0179, and Ta0020 were amplified from the genomic DNA. The other ITS sequences were derived from the NCBI database. Th3844: *T. harzianum* IOC-3844; Th0179: *T. harzianum* CBMAI-0179; Ta0020: *T. atroviride* CBMAI-0020.

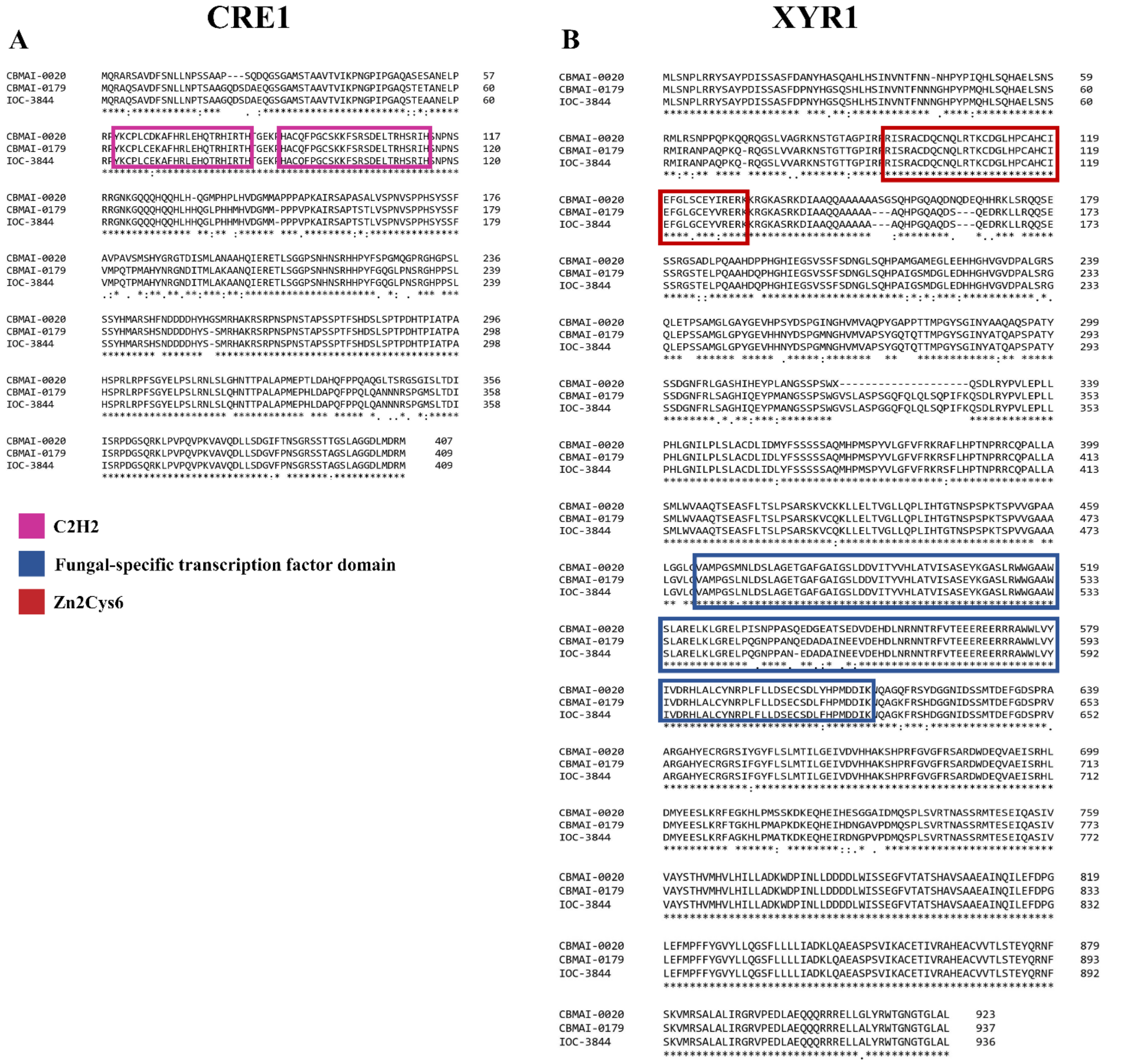


**Supplementary Material 1: Supplementary Figure 2. Sequence alignments of CRE1 and XYR1 homologs from Th3844, Th0179, and Ta0020.** CRE1 and XYR1 protein sequences of the studied *Trichoderma* spp. were employed for the alignment using Multiple Sequence Alignment of Clustal Omega software (Sievers et al., 2011). IOC-3844: *T. harzianum* IOC-3844; CBMAI-0179: *T. harzianum* CBMAI-0179; CBMAI-0020: *T. atroviride* CBMAI-0020. An * (asterisk) indicates positions that have a single, fully conserved residue.

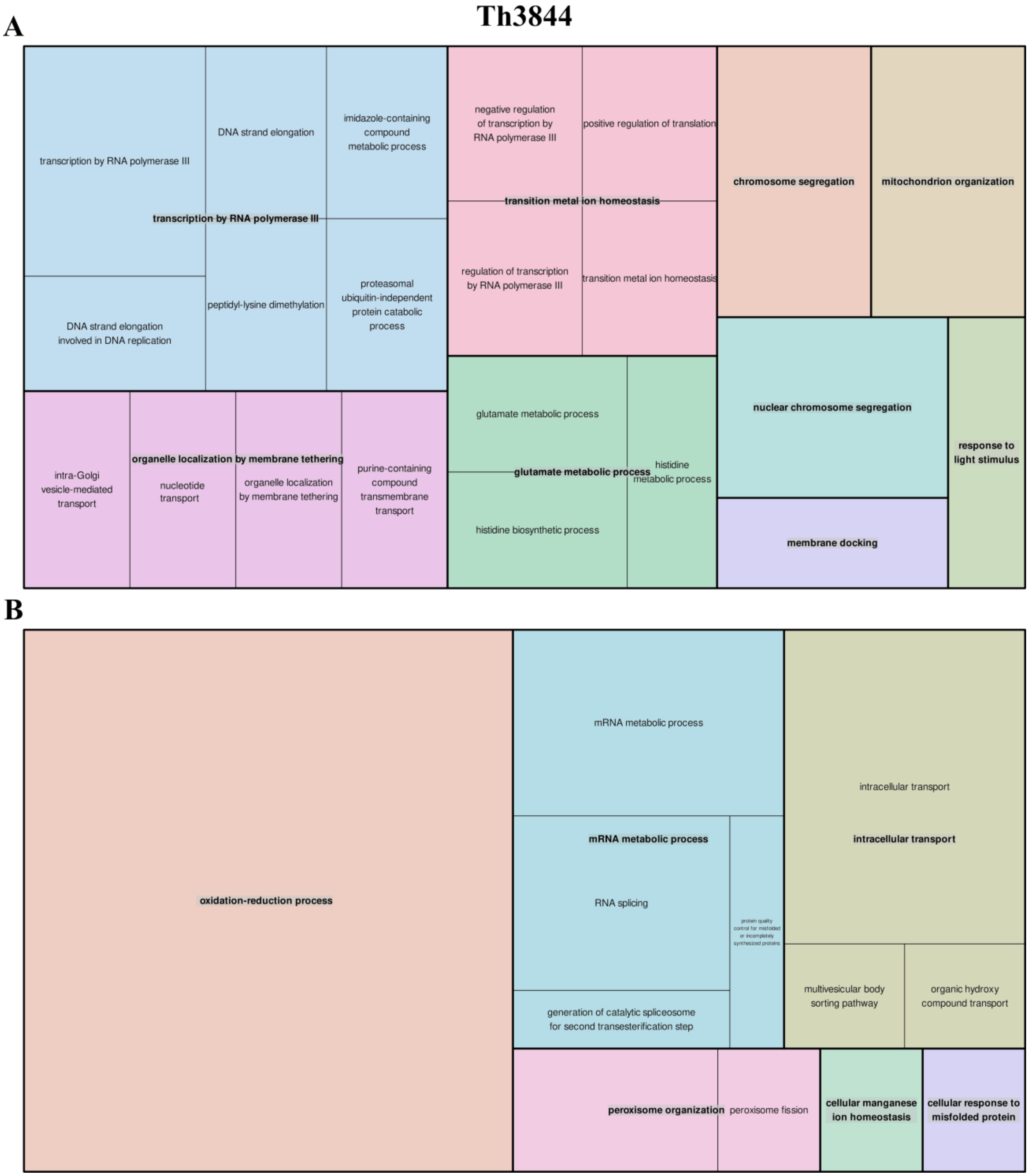


**Supplementary Material 1: Supplementary Figure 3. Treemap plotted based on enriched GO categories in the selected groups of Th3844.** GO analysis of all transcripts in the (A) *cre1* group and (B) *xyr1* group under the biological process category. Th3844: *T. harzianum* IOC-3844.

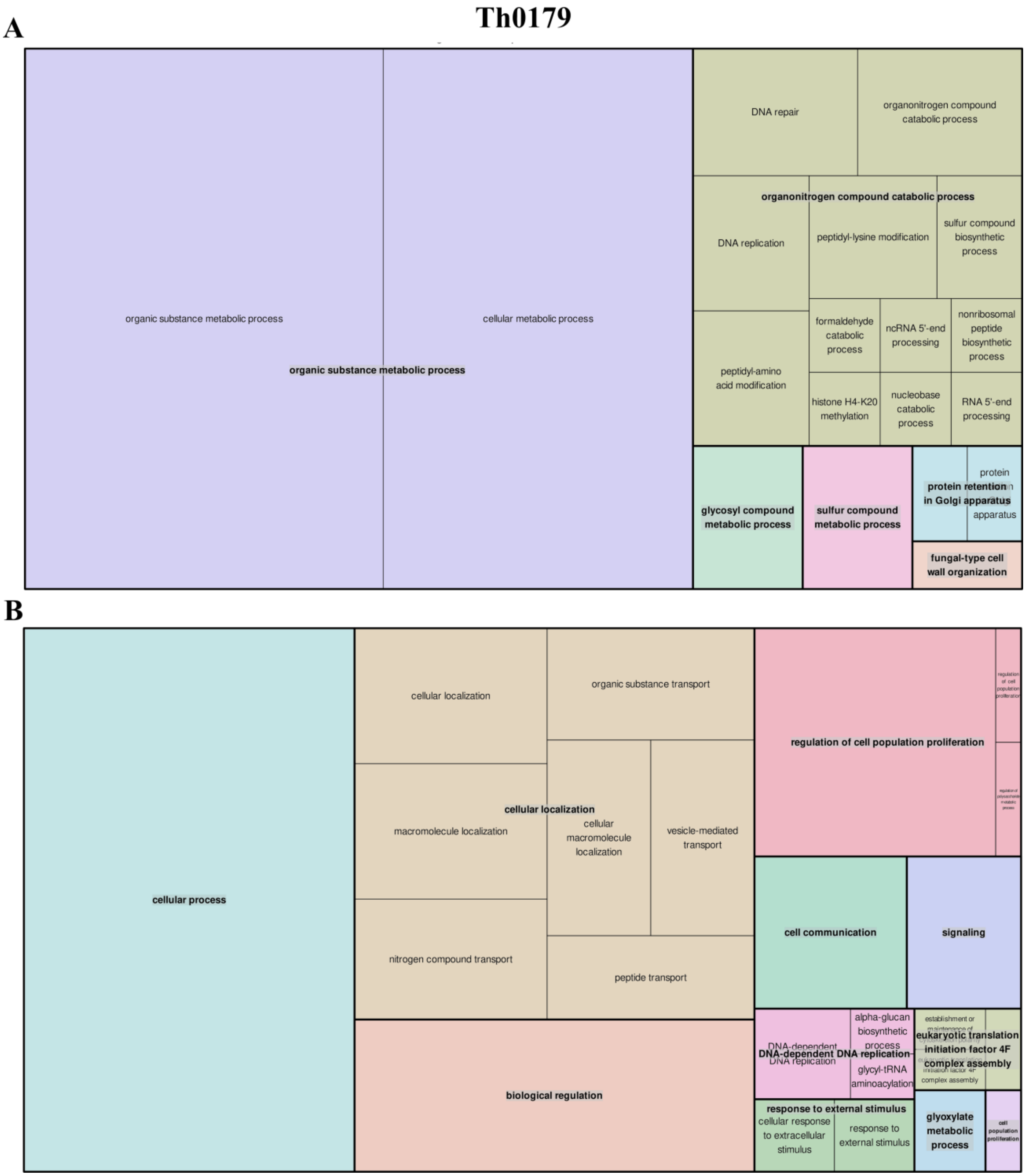


**Supplementary Material 1: Supplementary Figure 4. Treemap plotted based on enriched GO categories in the selected groups of Th0179.** GO analysis of all transcripts in the (A) *cre1* group and (B) *xyr1* group under the biological process category. Th0179: *T. harzianum* CBMAI-0179.

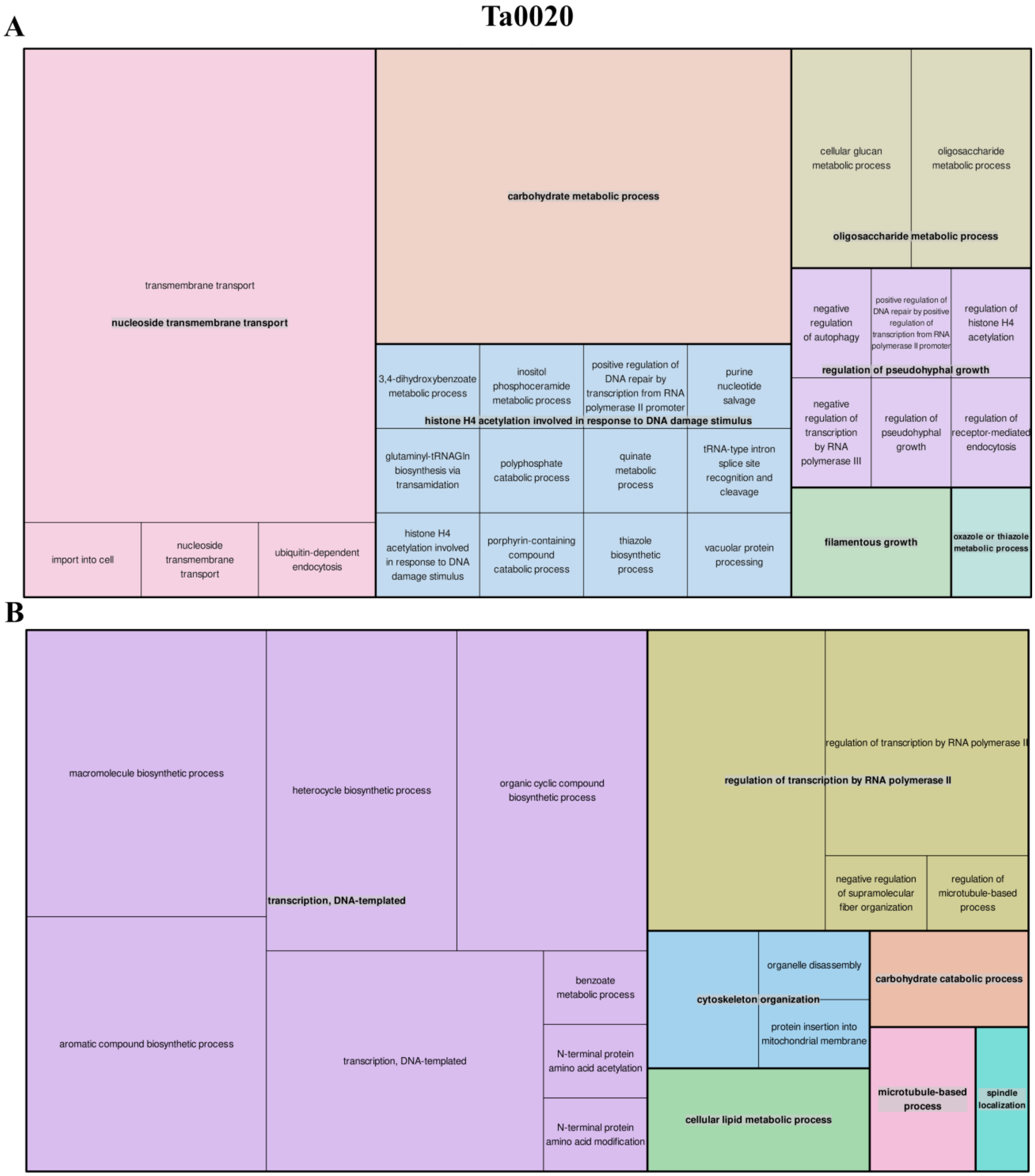


**Supplementary Material 1: Supplementary Figure 5. Treemap plotted based on enriched GO categories in the selected groups of Ta0020.** GO analysis of all transcripts in the (A) *cre1* group and (B) *xyr1* group under the biological process category. Ta0020: *T. atroviride* CBMAI-0020.

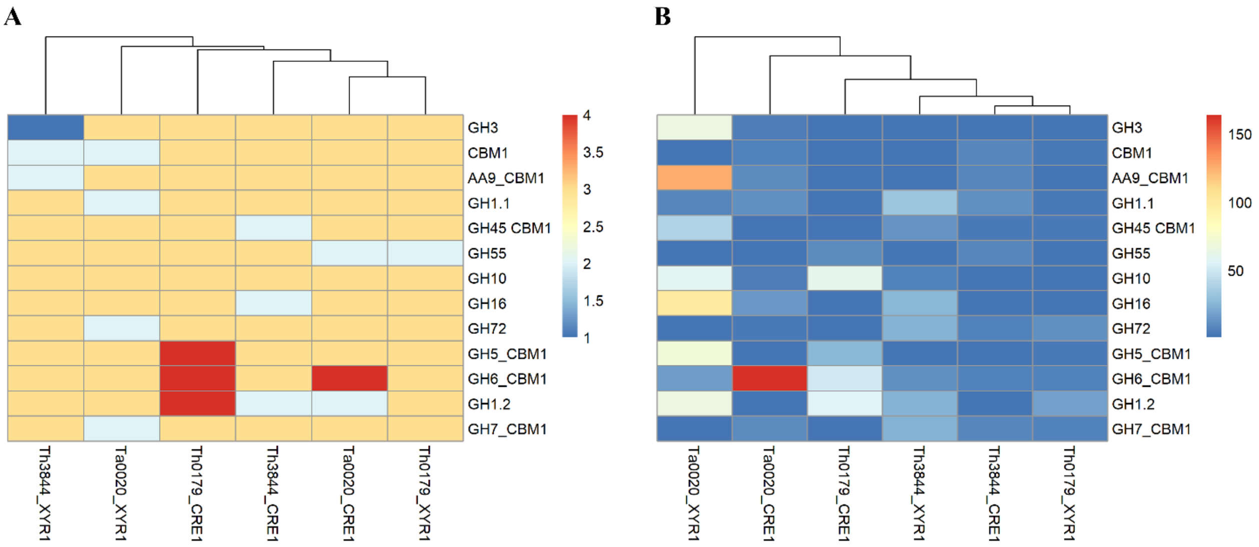


**Supplementary Material 1: Supplementary Figure 6. Heatmap plotted based on the shortest pathway between XYR1 or CRE1 and CAZymes in all evaluated *Trichoderma* spp.** Number of shortest pathways between XYR1 or CRE1 and the selected CAZymes (A) and number of possibilities of such events occurring (B). Th3844: *T. harzianum* IOC-3844; Th0179: *T. harzianum* CBMAI-0179; Ta0020: *T. atroviride* CBMAI-0020.

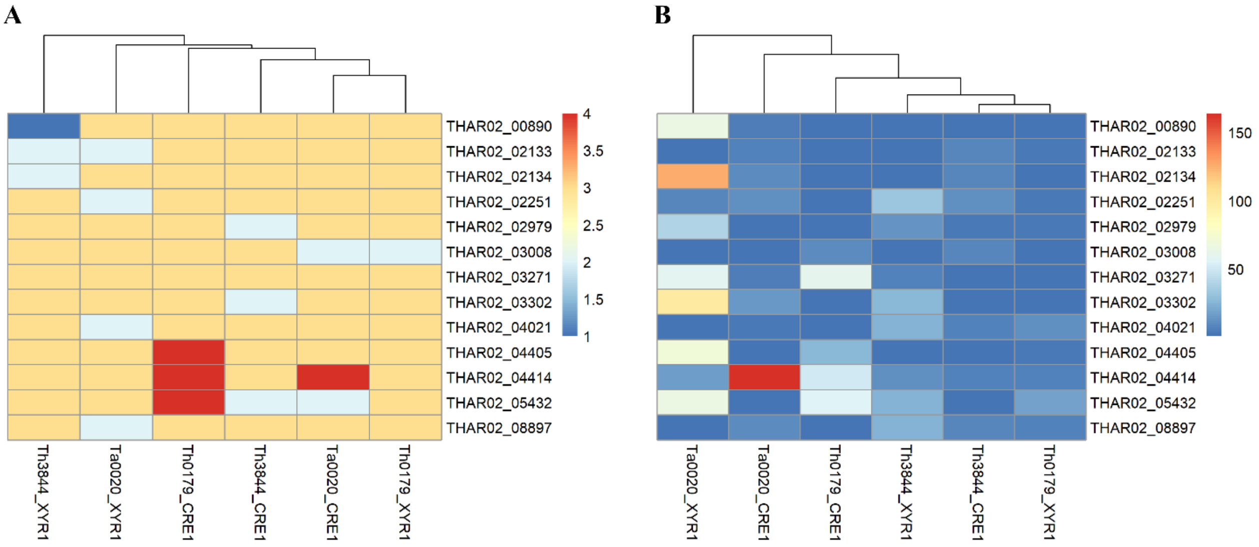


**Supplementary Material 1: Supplementary Figure 7. Heatmap plotted based on the shortest pathway between XYR1 or CRE1 and CAZymes in all evaluated *Trichoderma* spp.** Number of shortest pathways between XYR1 or CRE1 and the selected CAZymes (A) and number of possibilities of such events occurring (B). The CAZymes used for the heatmap are represented by their ID. Th3844: *T. harzianum* IOC-3844; Th0179: *T. harzianum* CBMAI-0179; Ta0020: *T. atroviride* CBMAI-0020.

**Supplementary Material 1: Supplementary Table 7.** Expression (in TPM) of *cre1* and *xyr1* in *Trichoderma* spp. under cellulose and glucose growth conditions.

| Strain | Gene ID | Protein product | FC values | p-value | Cellulose TPM | Glucose TPM | |
|---|---|---|---|---|---|---|---|
| **Th0179** | THAR02_10775 | KKO97123.1 | 1,041 | 0.271 | 906,063 | 854,791 | |
| **Th3844** | THAR02_10775 | KKO97123.1 | -1,136 | 0.035 | 519,282 | 598,932 | ***cre1*** |
| **Ta0020** | TRIATDRAFT_301116 | 013941427.1 | 2,096 | 0.000 | 1136,938 | 537,489 | |
| **Th0179** | THAR02_09260 | KKO98630.1 | -1,525 | 0.080 | 38,444 | 57,553 | |
| **Th3844** | THAR02_09260 | KKO98630.1 | 1,225 | 0.291 | 89,359 | 74,078 | ***xyr1*** |
| **Ta0020** | TRIATDRAFT_78601 | 013941705.1 | -1,649 | 0.056 | 23,978 | 39,190 | |

Expression profiles of *cre1* and *xyr1* among *Trichoderma* spp. were calculated in TPM. Genes with p-values ≤ 0.05 and fold changes ≥ 1.5 (upregulated) or ≤ -1.5 (downregulated) were considered differentially expressed. Th0179: *T. harzianum* CBMAI-0179; Th3844: *T. harzianum* IOC-3844; Ta0020: *T. atroviride* CBMAI-0020.
